# SOS-mediated prophage induction constrains resistance evolution to DNA-damaging antibiotics

**DOI:** 10.64898/2026.06.23.733845

**Authors:** Amy D. Zamora, Shreyas V. Pai, Kepler S. Mears, Fernando Rossine, Siân V. Owen, Michael Baym

## Abstract

Most naturally occurring bacteria are lysogens, encoding one or more temperate phages (prophages) integrated into their genome. As prophages are induced by the bacterial SOS response, DNA-damaging antibiotics can trigger SOS-mediated prophage induction, where prophages undergo lytic replication and lyse their host, even at sub-inhibitory concentrations. This prophage-antibiotic synergy therefore sensitizes lysogenic hosts to DNA-damaging antibiotics. However, the mechanism by which prophage-induced sensitization affects the evolution of resistance against these agents is unclear. Here we show that ciprofloxacin-resistant lysogens arise less frequently but exhibit higher levels of resistance following selection. Whole-genome sequencing showed that increased lysogen resistance arose from selection towards mutations in drug targets, efflux pathways, and stress response regulators that reduce antibiotic efficacy or alter SOS induction. Consistent with this result, resistant lysogens exhibited a dampened SOS response, suggesting that prophage induction imposes an additional selective filter on their hosts by eliminating mutants that experience sufficient DNA damage to activate the SOS response. By contrast, prophage carriage had no effect on sensitivity or resistance evolution for antibiotics where DNA damage occurs downstream of the primary mechanism of action. Together, these findings indicate that prophage induction acts as an evolutionary bottleneck that restricts many resistance trajectories while favoring the emergence of rarer, large-effect mutations, potentially accelerating the evolution of high-level resistance.

## Introduction

Antibiotics and bacteriophages are both major selective pressures which shape the evolution of bacteria. Notably, up to 75% of all sequenced bacterial genomes harbor prophages – bacteriophages that are integrated into a host chromosome.^1^ Consequently, many targets of antibiotic therapy are lysogens, where prophages are stably maintained until induced by host stress responses, particularly those triggered by DNA damage.^2^

Many antibiotics act by damaging DNA. These antibiotics select for mutations that decrease DNA damage by modifying binding targets or by reducing the intracellular concentration of antibiotic molecules. Fluoroquinolones, such as ciprofloxacin, primarily act by causing DNA gyrase to induce double-stranded chromosomal breaks.^3^ Common fluoroquinolone resistance mutations occur in characteristic region of DNA gyrase and topoisomerase IV, termed the quinolone resistance determining region (QRDR).^4,5^ Other resistance mutations are often found in efflux pump pathway genes, which upregulate pump activity to transport fluoroquinolones out of the cell or reduce porin permissibility.^4,6^

Bacteria can also survive mild DNA damage by SOS-mediated DNA repair.^7^ The accumulation of single-stranded DNA from this damage triggers the SOS response, where over 50 genes implicated in repair mechanisms are derepressed.^7,8^ Depending on the amount of DNA damage incurred, the SOS response can correct for double-stranded breaks and other genetic damage by expressing genes with functions ranging from excision repair to nucleotide synthesis via DNA polymerases.^7^

Although this conserved response has evolved to prevent death from DNA damage, it is also known to be a direct trigger for prophage induction, where prophages excise from the genome and resume their lytic cycle (Fig. 1A).^2,9^ This process uses host machinery to replicate and assemble virions, eventually killing the bacterial host through lysis where newly assembled progeny phages are released.^1,2,10^ This turns bacterial hosts’ attempt at repairing DNA damage into a paradoxically faster way to die.

**Figure 1.**
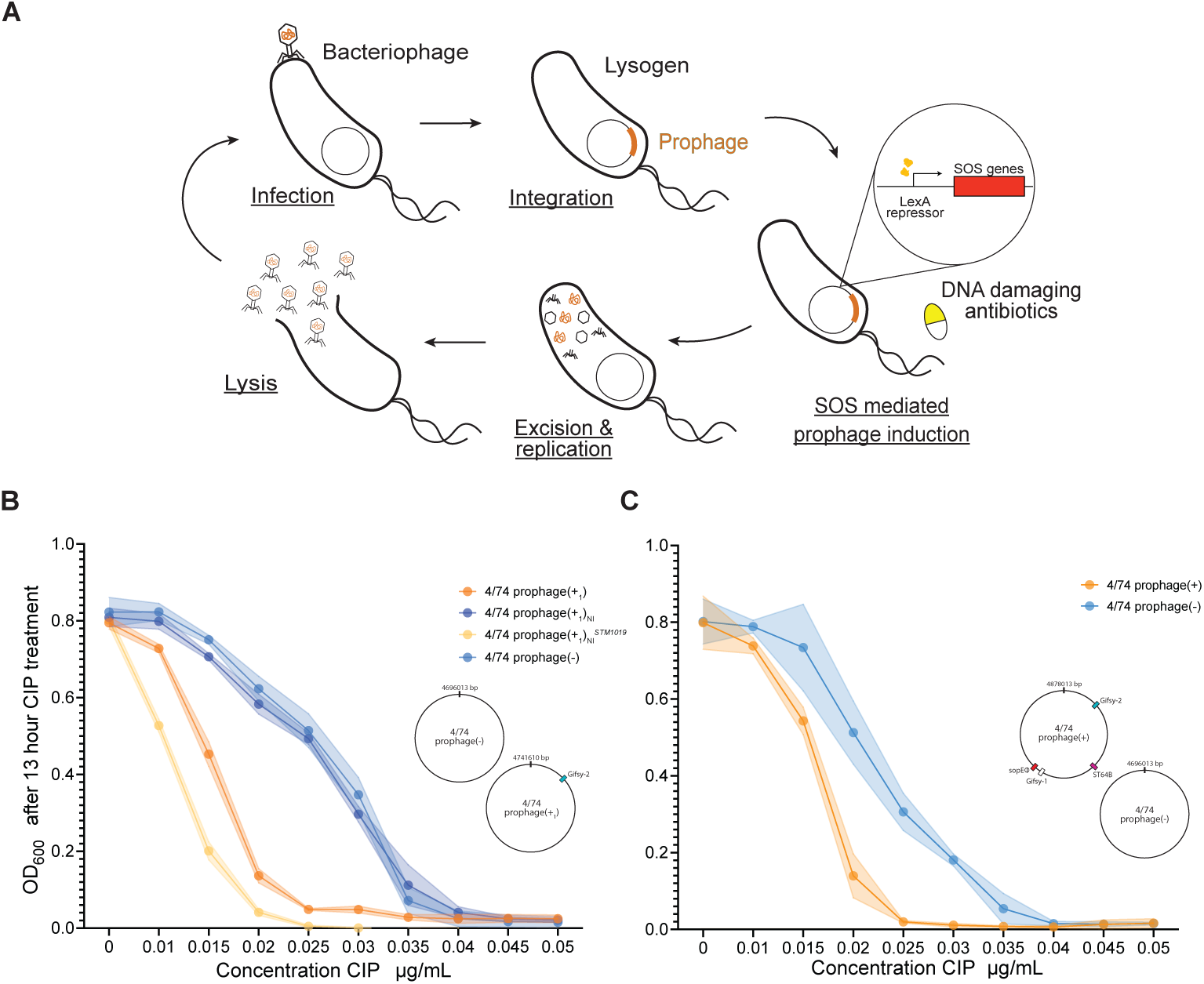
Prophage induction sensitizes *S.* Typhimurium to CIP. **A.** Model illustrating how DNA-damaging antibiotics can induce prophage excision and re-entry into the lytic cycle. **B.** OD_600_ values of single-lysogen 4/74 prophage(+_1_), a non-inducible (NI) knockout, a complemented knockout (*STM1019*), and a non-lysogen 4/74 prophage(-) following 13 h CIP treatment across a range of concentrations (x-axis) as a measure of sensitivity. Points represent mean ± SD of biological replicates. Survival of 4/74 prophage(+_1_) was significantly lower than both the NI knockout and 4/74 prophage(-) across intermediate CIP concentrations (multiple *t*-tests with FDR correction, adjusted *P* < 0.001). Complementation of the induction gene restored CIP sensitivity. **C.** Survival of the prophage(+) polylysogen and prophage(-) non-lysogen following 13 h CIP treatment. Prophage(+) survival was significantly lower than prophage(-) at intermediate CIP concentrations (multiple *t*-tests with FDR correction, adjusted *P* < 0.001). The sensitivity defect was comparable to that observed in the single-lysogen background.

Prophages have been observed to synergistically sensitize their hosts to various stresses, including antibiotics, reactive oxygen species, bacterial toxins/virulence factors, UV, and amino acid deprivation.^11–13^ Combination treatment with various antibiotics, including ciprofloxacin, and temperate phages has been shown to increase the sensitivity of lysogens through prophage induction at sub-inhibitory concentrations.^14–16^ For several of these stressors, this sensitization is hypothesized to work through prophage induction via the SOS response and avoidance of lysogeny upon infection.^14,17^ It has also been shown that sub-inhibitory concentrations of DNA-damaging antibiotics induce the SOS response and kill treated bacterial hosts through prophage induction, rather than antibiotic killing.^6,18,19^

In this study we explore how the SOS response, prophage induction, and fluoroquinolone antibiotics interact, asking whether the evolution of *de novo* resistance against DNA-damaging antibiotics differs depending on host prophage carriage.^20^ Using *Salmonella* Typhimurium as a model system, we examined population-level differences in ciprofloxacin resistance selection. We find that prophage-carrying hosts are more sensitive to ciprofloxacin as a direct consequence of SOS-mediated prophage induction. In turn, prophage carriage constrains resistance by skewing selection towards mutants with reduced SOS response activation, and consequently reduced prophage induction. This increased selection yields smaller but significantly more resistant populations enriched for high resistance mutations. Hence, we hypothesize that DNA-damaging selective pressures can accelerate the emergence of high-level resistance in lysogenic populations. However, this effect is not universal across SOS-inducing antibiotics, suggesting that prophage-antibiotic interactions depend on both the extent and timing of DNA damage.

## Results

### Increased sensitivity of lysogens to ciprofloxacin is caused by SOS-mediated prophage induction

To determine whether prophages sensitize *Salmonella* Typhimurium (*S.* Typhimurium) to ciprofloxacin (CIP), we measured growth in media supplemented with CIP for 3 different derivatives of strain 4/74: wild type (WT) containing 4 native prophages (prophage(+)), a 4/74 derivative with one intact prophage (prophage(+_1_)), and an isogenic prophage-cured derivative (prophage(-)) (Fig. S1).^21,22^ After 13 hour growth, the single lysogen prophage(+_1_) was 1.4-fold more sensitive to CIP than the cured prophage(-) (MIC 0.025μg/mL vs 0.035μg/mL, Fig. 1B). We observed the same increase in sensitivity of polylysogenic prophage(+) compared to prophage(-) (1.4x MIC, 0.025μg/mL vs 0.035μg/mL, Fig. 1C), showing that the number of resident prophages in a host does not affect the strength of sensitization.

SOS-mediated prophage induction further accounts for the increased antibiotic sensitivity of lysogens. A non-inducible single lysogen, 4/74 prophage(+_1_)_NI_, with its prophage’s antirepressor (*STM1019*)^23^ removed, exhibited CIP sensitivity comparable to that of prophage(-) (Fig. 1B), showing that prophage induction is directly responsible for enhanced CIP sensitivity in lysogens. Complementation of *STM1019 in trans* not only restored CIP sensitivity, but increased it, likely due to elevated antirepressor expression on a multicopy plasmid lowering the threshold for prophage induction.

### Resistant lysogens surviving Ciprofloxacin selection are rarer but more resistant

To investigate whether prophage-dependent differences in antibiotic sensitivity would affect the evolution of resistance, we selected spontaneous resistant mutants of polylysogenic 4/74 prophage(+) and 4/74 prophage(-) by plating on CIP after growth without antibiotics (Fig. 2A).

**Fig. 2.**
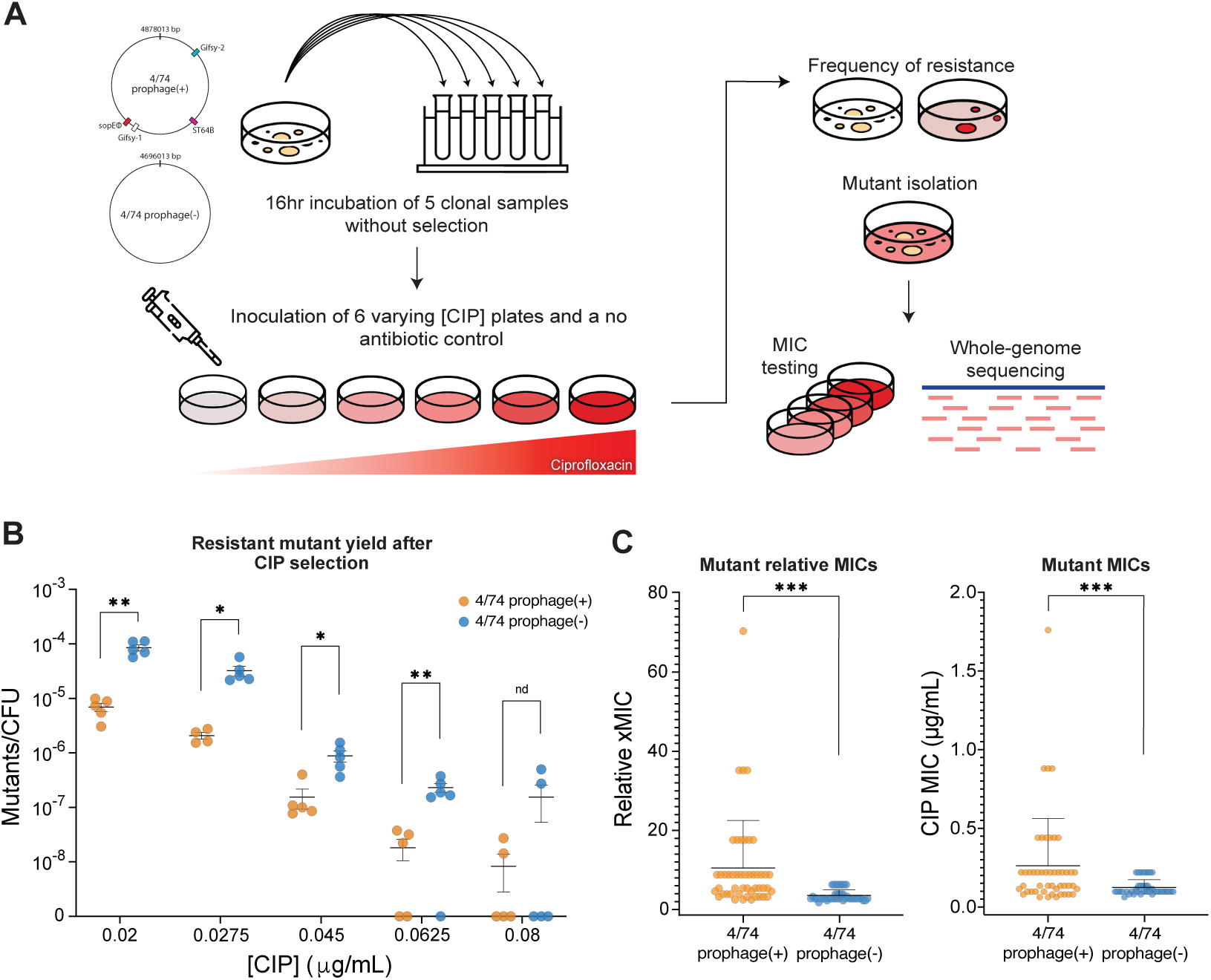
CIP-surviving lysogens are rarer, but more resistant. **A.** Experimental workflow for isolation of CIP-resistant mutants. Five independent clonal replicates of 4/74 prophage(+) and 4/74 prophage(-) were grown for 16 h in antibiotic-free liquid culture to avoid prophage induction, then plated on CIP gradient plates and antibiotic-free controls. After incubation, surviving colonies were counted and isolated for downstream phenotypic and genotypic analyses. **B**. Survival of lysogen and non-lysogen populations following CIP exposure across selective concentrations. Points represent individual plates from independent biological replicates. Survivor counts were normalized to paired no-antibiotic control plates. Prophage(+) lysogens showed significantly reduced survival relative to prophage(-) at all but one CIP concentration (multiple unpaired Mann–Whitney U tests). **C**. Relative and absolute MICs of resistant mutants isolated from panel B. Each point represents an individual mutant. Despite lower survival during selection, prophage(+)-derived mutants exhibited higher relative and absolute MIC values than prophage(-)-derived mutants (two-tailed Welch’s *t*-test on log2-transformed values, *P* < 0.001).

Ciprofloxacin-resistant mutants were 12.9x rarer across concentrations of CIP selection in the lysogenic background compared to the non-lysogen (Fig. 2B). Fluctuation assays for both ancestral strains showed that the baseline mutation rates of prophage(+) and prophage(-) strains were not measurably different (Fig S2B). This difference is likewise not explained by spontaneous induction, as seen in no-antibiotic growth conditions (Fig. 1B-C, S2A). Thus, as prophage carriage is the only difference between the strains, we conclude that the observed difference in survivor yield reflects SOS-mediated prophage induction killing a subset of mutants in the prophage(+) background.

Ciprofloxacin-resistant lysogens, while rarer, had a significantly higher average absolute MIC (0.263μg/mL vs 0.124μg/mL, Fig. 2C, *P*<0.001) and relative MIC (10.5x vs 3.5x, Fig. 2D, *P*<0.001) than prophage(-). While selection at the same concentration (0.0625 µg/mL) is a higher relative barrier for lysogens to overcome in resistance adaptation (2.5x MIC vs 1.8x MIC), this difference is far smaller than the observed differences in relative mutant MIC (Fig. 2C). Overall, antibiotic-resistant lysogens emerged less frequently; however, those that survived selection exhibited higher levels of resistance, suggesting that prophage-dependent differences in mutant selection promote larger adaptive steps.

### Prophage carriage alters the distribution of mutated pathways and clinically relevant CIP resistant mutations

We then asked how the identities of resistance mutations differed in the presence of inducible prophages. We isolated and sequenced 68 mutants (33 prophage(+) & 35 prophage(-)) from the 0.0625 µg/mL selective CIP concentration, as it is just above the threshold of “clinical resistance” for *Salmonella* (see Methods).^24^ As we cannot distinguish whether identical mutations in the same experiment are parallel events or clonal expansions of a single event, we report both the absolute number of mutations and an adjusted count of mutations with duplicates from the same replicate removed (“unique and independent” mutations, 54 of 68 absolute mutations, Fig. S4, Table S2).

We classified unique and independent mutations into affected pathways based on reported functions of mutated genes, with genes participating in shared resistance mechanisms or cellular processes assigned to the same class (see Methods). For these unique mutations, both prophage(+) and (-) yielded mutations in similar genes and gene pathways, but the overall distributions of the affected gene pathways were significantly different (Fig. 3, *P*<0.0001).

**Fig. 3.**
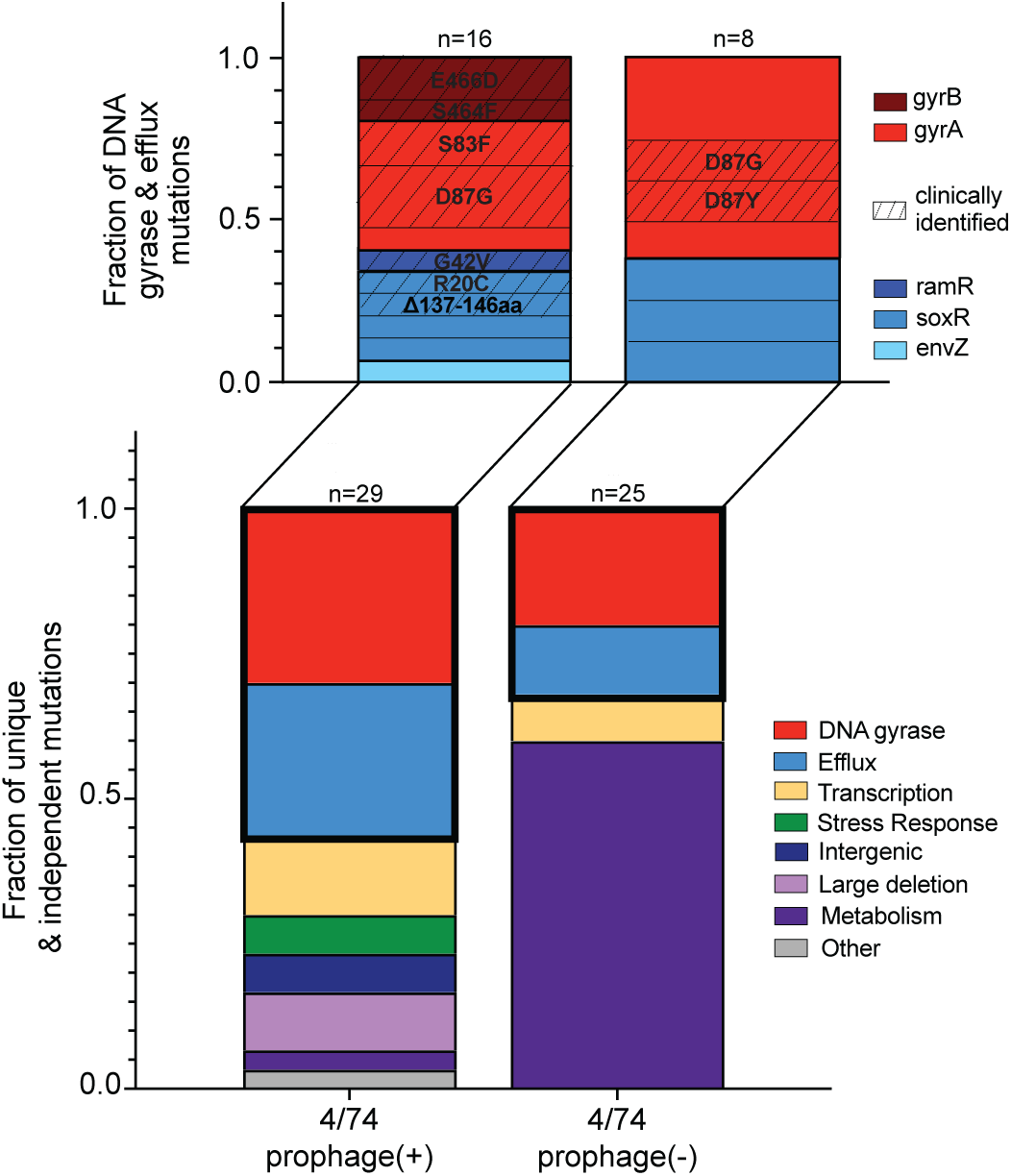
Mutated pathway distributions depend on prophage carriage. Bars show the distribution of pathways/categories represented by genes carrying resistance-conferring mutations among independent mutants from 4/74 prophage(+) and prophage(-) backgrounds. Each colored segment indicates the fraction of total unique mutations assigned to a given host function pathway. Pathway distributions differed significantly between ancestral backgrounds (permutation chi-square test, 10,000 label shuffles, P<0.0001). Canonical quinolone resistance pathways (DNA gyrase and efflux pump) were mutated in both backgrounds. These categories comprised a larger fraction of mutations in prophage(+), but this enrichment was not significant (Monte Carlo permutation test; Fisher’s exact test, n.s.). The expanded inset shows the composition of mutations within the DNA gyrase and efflux pump categories, with each subsection representing a specific mutation and its fraction of total unique mutants. Mutations previously associated with clinical resistance are indicated by striping, with amino acid substitutions labeled over top.

We overall observed more mutations in DNA gyrase and efflux pathways in the lysogenic background (Fig. 3, Fig. S5). These two well-established quinolone resistance pathways constituted 57% of the unique and independent prophage(+) CIP-resistant mutants compared with 32% in the prophage(-) background. Additionally, many of the DNA gyrase mutations overlapped with previously described fluoroquinolone resistance mutations associated with high level resistance in clinical isolates (Fig. 3, Table S3).^5,6,25–27^ While less well characterized, several, efflux-associated genes were also mutated at residues or within regions previously implicated in elevated, clinical resistance (Fig. 3, Table S3).^28–31^ Although the number of mutations in these genes were different, resistance in these specific pathways was expected, as DNA gyrase is the direct target of CIP and the affected efflux genes are regulators of the AcrAB system, which exports CIP into and out of the cell.^4,5,27,28,32^ Thus, in addition to selecting for mutations conferring greater resistance, prophage carriage appears to enrich for mutations on the path to very high-level resistance.

The largest observed difference in distributions was that prophage(-) resistant mutants were significantly enriched for metabolic pathway-related mutations (Fig. 3, *P* < 0.0001). In particular, metabolism-related mutations in the prophage(-) background were only observed in a single gene, *purB*, a purine biosynthesis gene.^33,34^ No point mutations were detected in *purB* in lysogens. Instead, the only metabolic mutation observed was a single SNP in *ribD,* a riboflavin biosynthesis gene.^35^

Several prophage(+) and prophage(-) resistant mutants carrying mutations in *gyrA*, *soxR, ramR,* and *purB,* also contained secondary mutations. As many of these mutations did not have functional ties to CIP resistance and were only seen in combination with other resistance-conferring mutations, they were considered to be putative compensatory mutations (Tables S2-3).

Several pathways were uniquely mutated in lysogenic mutants. We identified two mutations occurring in *clpP*, a stress response-associated protease,^36,37^ as well as two mutations in the intergenic region between putative transcriptional regulator *yiaG* and major cold-shock protein *cspA*,^38^ occurring across parallel lysogen replicates. Similarly, we observed large-scale (>2kb) deletions in three parallel lysogen replicates. Two of these deletions overlapped with each other, despite being in separate populations. Although these deletions did encompass genes which confer CIP resistance on their own (*envF, icdA, purB, hnr,* etc., Table S2),^33,39,40^ their size made it impossible to assign a single gene as resistance determining.

Higher frequencies of DNA gyrase and efflux mutations, together with the repeated occurrence of *clpP* stress response mutants in the prophage(+) background, suggested that survival during CIP selection was determined not only by resistance level, but also by the extent to which SOS-mediated prophage induction occurs. Notably, *clpP* mutations have been previously linked to stabilization of the SOS repressor LexA.^36,37^ Because several of our active prophages require LexA cleavage for induction,^20,23^ we hypothesized that survival of CIP-resistant lysogens depends on both the ability of CIP to generate DNA damage and the subsequent induction of the SOS response in selection. Under this model, prophage carriage does not alter which resistance-conferring mutations arise, but instead imposes an additional selective filter, eliminating cells that experience sufficient DNA damage to trigger SOS-dependent prophage induction. Consequently, surviving CIP-resistant lysogens are expected to more frequently harbor mutations that reduce DNA damage or dampen the SOS response, thereby preventing prophage induction.

### CIP-resistant lysogens exhibit lower levels of SOS induction

To test whether lysogens reduce DNA damage or SOS response to survive CIP exposure, we sought to assess the level of SOS induction experienced by mutants in both backgrounds. We expected that lysogenic resistant mutants overall have a lower threshold of viable SOS induction, as high SOS inducing survivors would be eliminated through prophage induction (Fig. 4A). We constructed reporters of SOS response induction in our evolved mutants by fusing the promoter of the SOS regulatory gene, *recA,* to GFP (precA-GFP) and engineering it into the chromosome of 19 of our CIP resistant mutants from both prophage(+) and prophage(-) backgrounds as well as our ancestor strains. We measured SOS induction via *recA* expression with and without exposure to sub-inhibitory concentrations of CIP (mutants at selection concentration 0.0625 µg/mL and ancestors at 0.01 µg/mL). Resistant prophage(+) mutants had broadly lower, but non-zero, SOS induction compared to prophage(-) mutants upon treatment with CIP (Fig. 4B, 1.36x lower average, *P* = 0.0114). The highest *recA*-expressing, and therefore SOS-inducing, samples were *purB* mutants for prophage(-), and a mix of efflux and stress response mutants for prophage(+) (Fig. S7, Table S3). Thus, SOS-mediated prophage induction selects against mutants with SOS levels above the phage induction threshold.

**Fig. 4.**
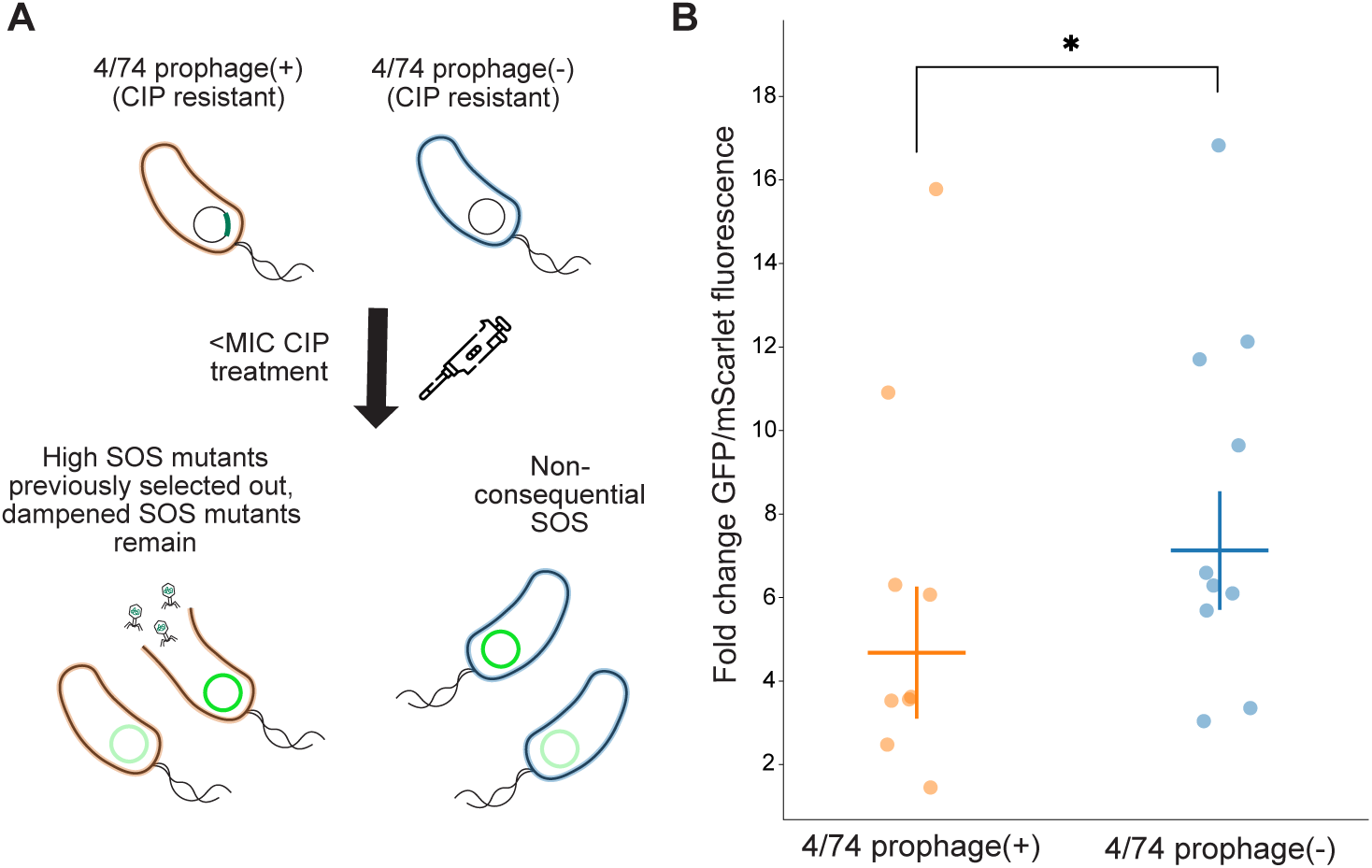
CIP-resistant lysogens exhibit a dampened SOS response. **A.** Model predicting that CIP-resistant 4/74 prophage(+) lysogens are enriched for mutations that reduce SOS induction and thereby avoid prophage induction during selection. Under sub-inhibitory CIP exposure, prophage(+) mutants are therefore expected to show lower SOS activation than prophage(-) mutants. SOS response was measured using a GFP reporter under control of the *recA* promoter. **B**. Fold change in GFP reporter signal (normalized by constitutive mScarlet fluorescence for cell size) between −CIP and +CIP conditions measured by flow cytometry. Each point represents a single mutant’s precA-GFP expression as a mean of three biological replicates. Prophage(+) mutants showed lower induction than prophage(-) mutants (Mann–Whitney U test, *P* = 0.0114).

We noticed a single prophage(+) mutant (s111, Table S3) had a much higher level of reporter expression than the rest of the lysogenic mutants. This strain contained a premature stop codon in clpP, which has been observed to stabilize LexA and prevent SOS-mediated prophage induction.^23,37^ Because LexA stabilization, or impaired LexA cleavage, occurs downstream of increased *recA* expression, our precA-based reporter may not accurately report completion of the SOS response in *clpP* or *lexA* mutants. Thus, s111 may still show increased expression after CIP treatment while failing to activate downstream SOS functions required for prophage induction. We therefore consider this fluorescent reporter unreliable for s111 and mutants with similarly suppressed SOS responses.

Resistant lysogens exhibit a dampened SOS response following CIP treatment. Mutants that either avoid SOS induction or reduce CIP-induced DNA damage survive in the lysogenic background. In contrast, prophage(-) mutants face no prophage-mediated cost for sustained SOS induction, allowing survival of a broader population of CIP-resistant mutants. Therefore, resistant lysogenic mutants with strong SOS induction are counter-selected by prophage-induced host death.

### Prophage-dependent differences in sensitization and resistance selection are not universal across SOS-inducing antibiotics

To determine whether the effects of prophage carriage extended beyond CIP, we examined host responses to an additional DNA-damaging antibiotic, mitomycin C (MitC),^41^ and a non-DNA-damaging antibiotic, trimethoprim (TMP).

MitC is a well-characterized DNA-damaging agent that triggers the SOS response and subsequent prophage induction (Fig. S3).^41^ We tested whether lysogens experienced increased sensitization to MitC as a consequence of SOS-mediated prophage induction. We challenged 4/74 prophage(+_1_), 4/74 prophage(+_1_)_NI_, 4/74 prophage(+_1_)_NI_*^STM^*^1019^, and 4/74 prophage(-) strains with MitC. Lysogenic prophage(+_1_) displayed increased sensitivity when compared to its prophage-free counterparts (Fig. 5A).

**Fig. 5.**
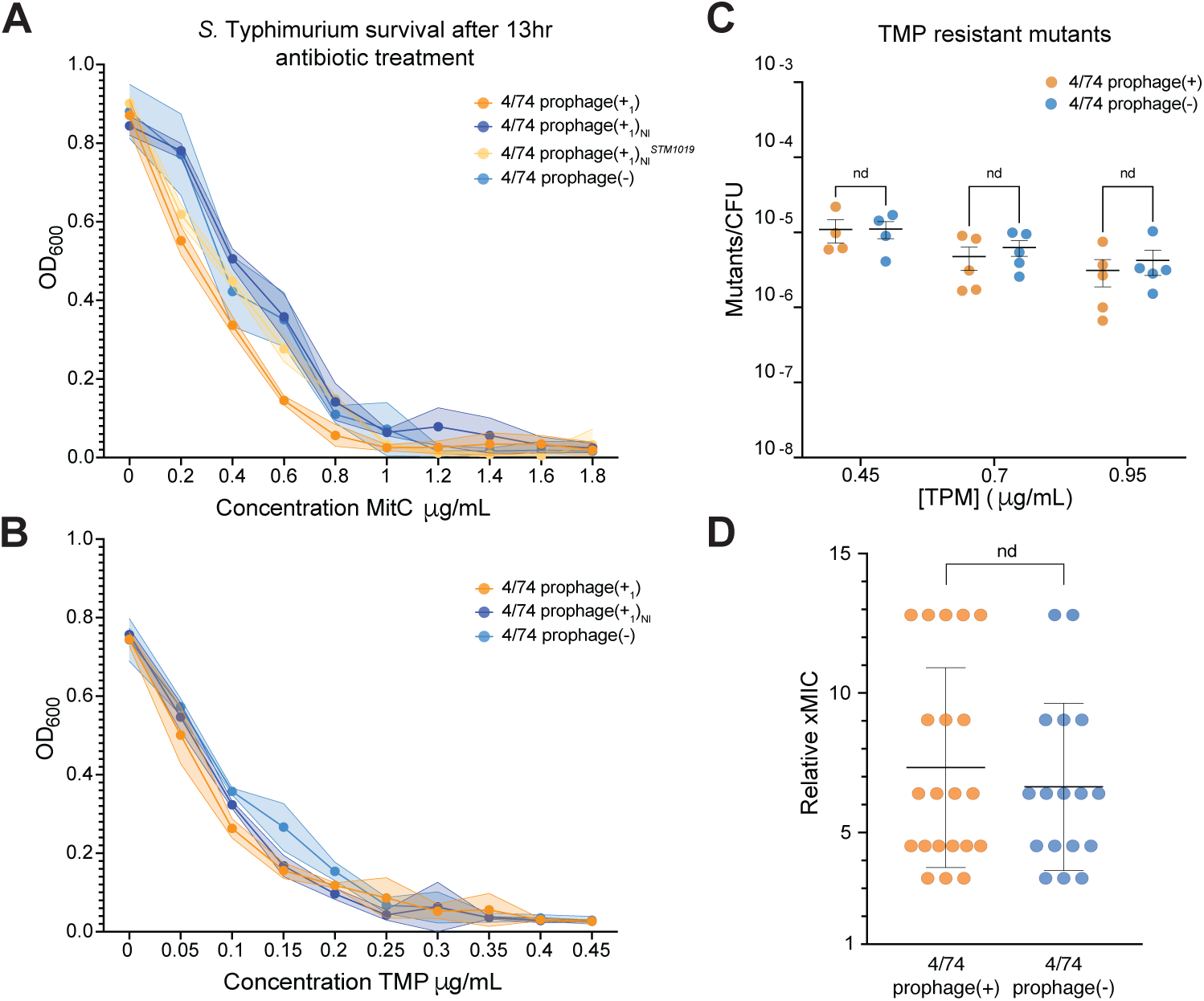
Prophage-dependent sensitivity and resistance selection differences vary by antibiotic. **A**. MitC sensitivity of single lysogen 4/74 prophage(+_1_), its non-inducible (NI) knockout, the complemented NI strain, and 4/74 prophage(-) measured by OD600 after 13 h of growth in a range of drug concentrations (x-axis). Lysogens showed significantly increased sensitivity at low-to-intermediate MitC concentrations relative to prophage-free or NI counterparts (prophage(+) vs prophage(-), *P* < 0.0001; prophage(+_1_) vs prophage(+_1_)_NI_/prophage(-), *P* < 0.05; multiple *t*-tests with FDR correction). Although complementation increased sensitivity relative to the NI knockout (*P* < 0.05), it did not fully restore the original lysogenic phenotype. **B**. Sensitivity testing as in panel A, but with TMP. Differences in sensitivity were observed at middle TMP concentrations between prophage(+_1_) and prophage(-) (*P* < 0.01); however, the induction knockout did not increase host sensitivity as observed with MitC and CIP, suggesting prophage induction is not the primary driver of minimal TMP sensitivity differences. **C**. Survival of TMP-resistant mutants following 16 h non-selective growth and subsequent selective plating. Each point represents an independent clonal replicate. **D.** Relative TMP MICs of surviving isolates. Each point represents an individual isolated resistant mutant. No significant differences were observed in mutant survival or relative TMP MIC values between backgrounds (multiple unpaired Mann–Whitney U tests; two-tailed Welch’s *t*-test on log2-transformed values, n.s.).

While the MitC sensitivity of the prophage(+_1_)_NI_ strain was decreased to the level of prophage(-), *STM1019* complementation did not fully restore sensitivity. While the polylysogenic prophage(+) strain also showed increased MitC sensitivity, it was at a much higher magnitude than the single lysogen (Fig. S9). This suggested that although prophage induction contributes to MitC sensitization, Gifsy-2 induction alone is insufficient to explain the heightened sensitivity observed in the prophage(+) background.

In contrast, TMP, despite inducing SOS, does not show similar synergy with prophage carriage. Although TMP does not directly damage DNA, secondary replication stress can induce low-level SOS activation via reactive oxygen species (Fig. S3).^42,43^ We observed that although TMP triggers the SOS response, sensitivity increases were minimal, regardless of whether strains were single lysogens (prophage(+_1_)) or polylysogens (prophage(+)) (Fig. 5B, Fig. S9). These minimal increases in sensitivity appeared to be prophage independent, as we see our induction knockout does not reduce this sensitivity. These findings show that although prophages sensitize hosts to direct DNA-damaging antibiotics such as ciprofloxacin and MitC, the extent of sensitization depends on the mechanism and magnitude of DNA damage generated by the antibiotic, likely through effects on SOS-mediated prophage induction. Indeed, we observed substantially lower SOS induction from TMP compared to direct DNA damaging agents CIP and MitC (Fig. S3), suggesting that a threshold level of DNA damage or SOS induction may be required to produce the prophage-dependent phenotypes.

With these observations, we wanted to understand if these minor lysogen increases in sensitivity affected TMP resistance trajectories during selection. In contrast to CIP, we observed no difference in mutant yield across backgrounds nor was there a difference in the mutant average relative MIC between prophage(+) and prophage(-) isolated mutants (see Methods) following TMP selection (Fig. 5C). This suggested that, despite causing secondary DNA damage and weak SOS activation, TMP fails to elicit substantial prophage induction, if any, and therefore does not alter resistance selection trajectories.

## Discussion

We showed that prophage carriage increases the efficacy of direct DNA-damaging antibiotics CIP and MitC against lysogens by triggering prophage induction at otherwise sub-inhibitory concentrations. We additionally find that CIP selection in lysogenic strains yields fewer, more resistant mutants with a dampened SOS response. However, this effect is not universal for SOS-inducing antibiotics, as observed with TMP. This suggests a model for prophage-antibiotic interaction, where the synergistic killing of the host is dependent on the SOS response directly ameliorating the antibiotic-induced damage in non-lysogens (Fig. 6A). The difference in antibiotic sensitivity with and without the phage is then reflected in the surviving resistance mutations, with the prophage restricting survivor mutations to those that directly decrease the effect of the drug and do no necessitate or increase SOS mediated repair (Fig. 6B).

**Fig. 6.**
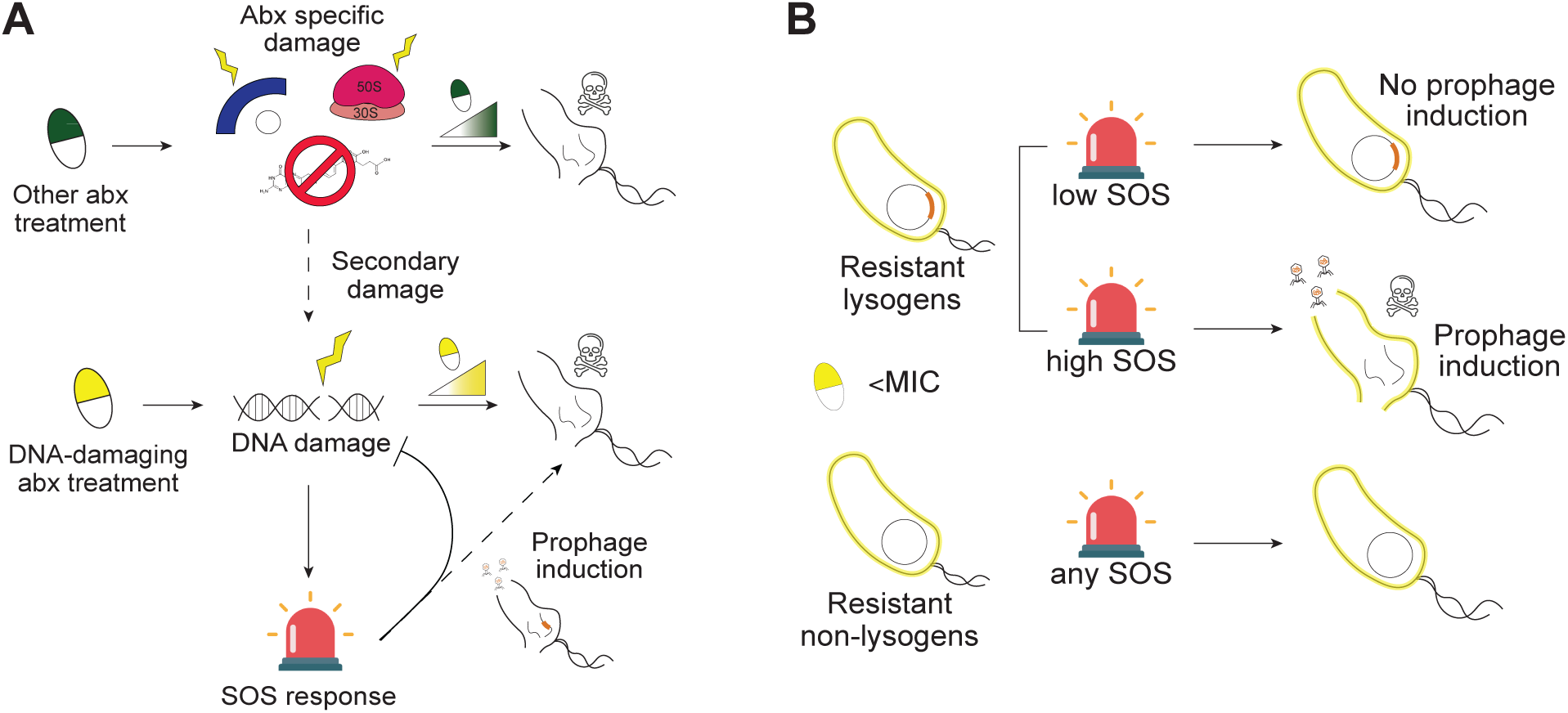
A model for prophage modulation of antibiotic sensitivity and selection. **A.** DNA-damaging antibiotics induce the SOS response at sub-lethal concentrations, allowing repair in non-lysogens but triggering prophage induction and host death in lysogens. Other antibiotic classes can generate secondary DNA damage at sub-inhibitory concentrations, activating SOS and prophage induction depending on mechanism. In these cases (e.g. TMP), as SOS activation does not address primary antibiotic damage in non-lysogens and may not activate the SOS enough to trigger prophages in lysogens, there is no difference in sensitivity or resistance selection. Thus, the mechanism of action for each antibiotic (including secondary effects) will ultimately determine this relationship. **B**. During DNA-damaging antibiotic resistance selection, prophage induction increases selection in lysogens: mutants that fail to dampen SOS or reduce DNA damage (and avoid prophage induction) are eliminated. In non-lysogens, mutants survive regardless of resistance mechanism.

This phenomenon likely extends to other DNA-damaging stressors whose effects are partially mitigated by SOS-dependent repair pathways, including UV irradiation and bacterial toxins. Indeed, DNA-damaging compounds produced during interbacterial competition have been shown to trigger prophage induction in neighboring lysogens through SOS activation.^13^ Although resistance mechanisms to these stressors may differ, any adaptation that allows robust SOS induction would still result in prophage-stress synergy.

In contrast, other indirect or non-DNA-damaging antibiotics are unlikely to follow this pattern. With TMP, we observed minimal prophage-dependent differences in sensitivity and no differences in resistance selection. This is consistent with previous literature showing no synergy between TMP and prophage induction without the addition of temperate phages above MOI of 1.^15^ Although TMP damages DNA, likely as a downstream effect of thymine starvation,^43^ its primary mechanism of action is blocking DNA synthesis.^44^ This could affect the ability of prophages to replicate even if induced, preventing the expression of genes required for host lysis, including holins.^2^ Alternatively, indirect DNA-damaging antibiotics may not cause enough downstream DNA damage (and in turn SOS response) to trigger prophages alone, yielding a similar effect to non-DNA-damaging antibiotics.

There are also prophages for which this phenomenon does not apply. Recent work not only shows that selective, SOS-independent prophage induction is possible,^45,46^ but also that defective prophages can yield incomplete prophage induction that avoids host lysis altogether.^47,48^ This would be expected to interact with resistance as there would be no lethal repercussions of prophage induction.

Additionally, we saw that the SOS response was dampened, not ablated, in lysogenic, CIP-resistant mutants. This implies that the level of SOS response (and therefore the amount of DNA damage incurred)^5^ by specific mutants during selection is vital in avoiding SOS-mediated prophage induction. Over longer evolutionary timescales, exposure to DNA-damaging stresses may therefore act as a pressure to tune the sensitivity of resident prophages’ lytic switches.^49^

The specific paths to resistance seen in lysogens mimic clinical observations. In DNA gyrase, a majority of the mutations conferring CIP resistance in lysogens were clinically relevant and had been previously shown to confer high-level resistance.^5,24,26^ Of these, we observed mutations in *gyrB* (the secondary subunit of DNA gyrase) exclusively in lysogens. We also observed mutations in efflux-related genes (*ramR* & *soxR* in the AcrAB system), specifically in residues implicated in clinical high-level resistance.^28–32,50^ Increased selection for these high-level resistance trajectories in lysogens suggests that prophage carriage may influence the long-term evolution of antibiotic resistance. Despite a smaller resistant mutant yield, selection of these trajectories in lysogens takes larger steps in resistance evolution. In environments with significant clonal interference, the rarity of high-resistance mutations in non-lysogens would compound, decreasing the frequency of mutants with multiple strong mutations required for high-level resistance. Thus, while prophages constrain immediate pathways to resistance, our results show their potential to paradoxically accelerate high-level resistance evolution.

## Methods

### Bacterial Strains and Culture Conditions

A complete list of strains and plasmids used in this study is provided in Table 1. Primary experiments were performed in *Salmonella enterica* serovar Typhimurium strain 4/74 wild type (4/74 prophage(+); accession CP002487), prophage-cured 4/74ΔΦ [ΔsopEΦ, ΔST64B, ΔGifsy-1, ΔGifsy-2] (4/74 prophage(-)), and their evolved or genetically modified derivatives. 4/74 prophage(+) contains four naturally integrated prophages (sopEΦ, ST64B, Gifsy-1, and Gifsy-2). 4/74 prophage(-) was previously generated by scarless genome editing^11,38^ The strain is isogenic to 4/74 prophage(+) except for silent mutations at positions 2,900,561; 4,693,754 (*iolC*); and 1,870,861 (*minE*). Additional genetic modifications were generated using the same scarless editing approach. *Salmonella enterica* serovar Typhimurium strain D23580ΔΦ *waaG:aph* was used exclusively as an indicator strain for plaque assays.

**Table 1:**
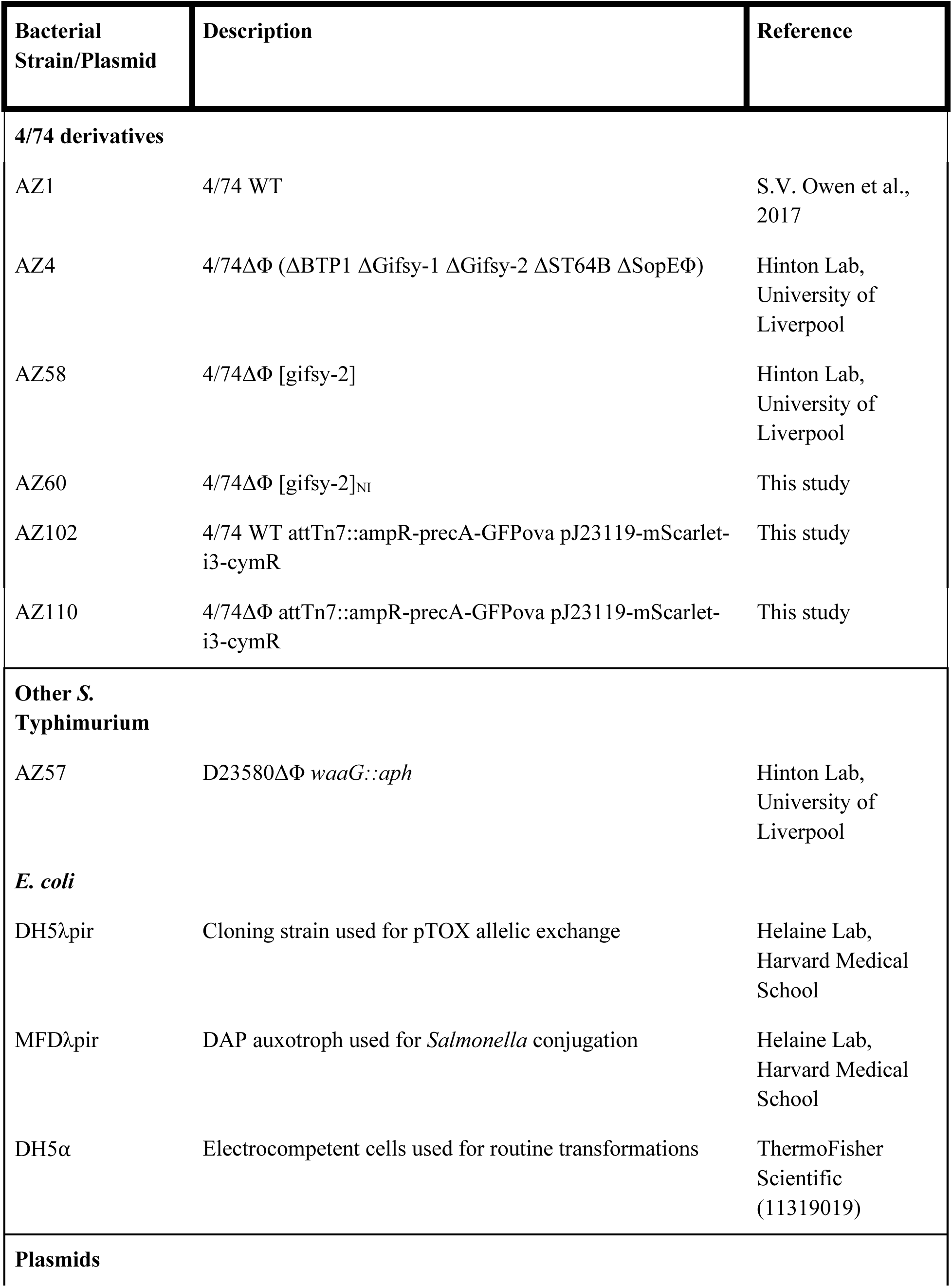
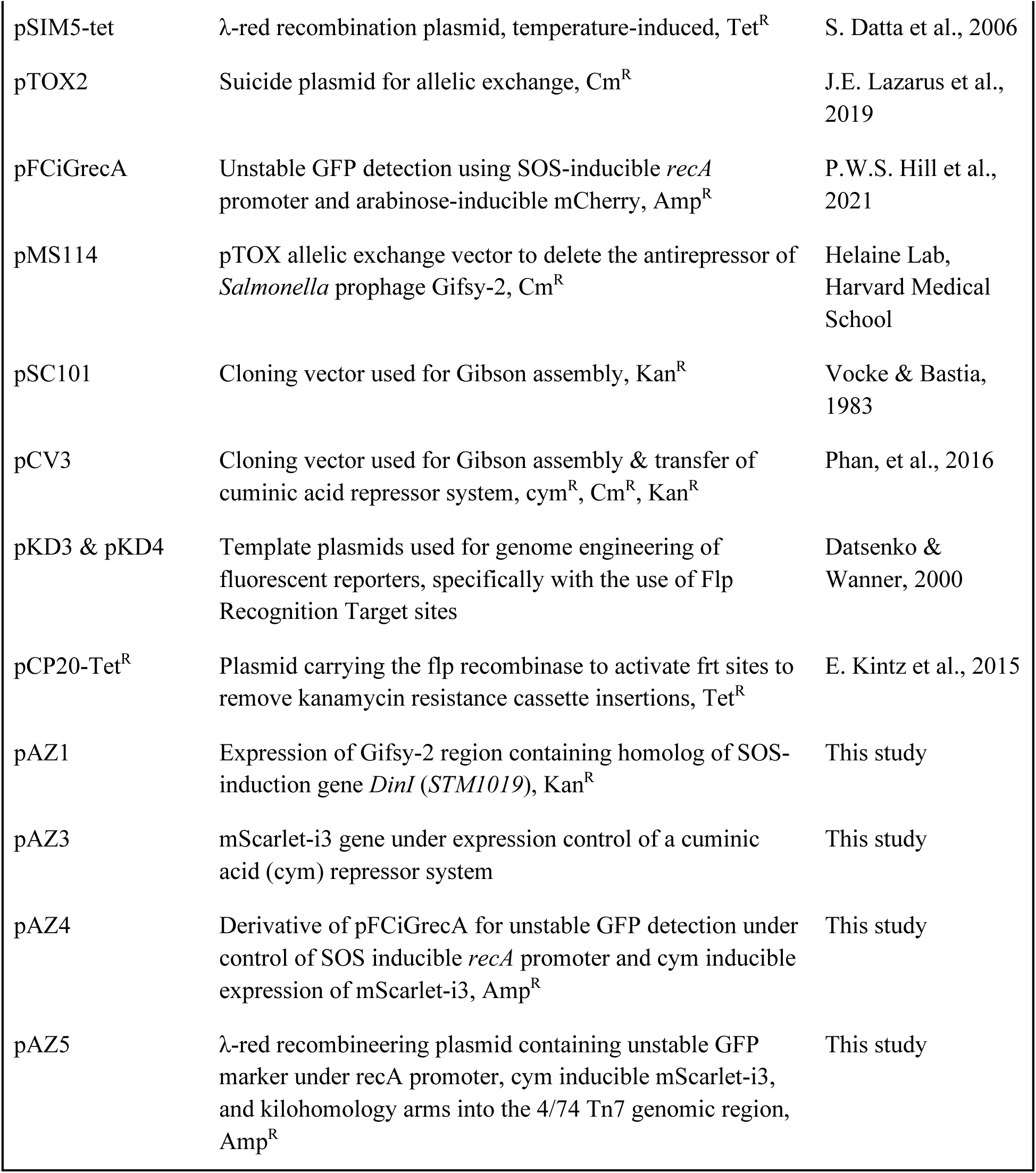
Bacterial strains and plasmids used in this study.

Unless otherwise stated, strains were cultured in LB Lennox (Research Products International L24066) containing 10 g/L tryptone, 5 g/L yeast extract, and 5 g/L NaCl (pH 7.0). Where indicated, LB or LB agar (2%) was supplemented with ampicillin (100 µg/mL), carbenicillin (100 or 500 µg/mL), chloramphenicol (34 µg/mL), kanamycin (50 µg/mL), or tetracycline (20 µg/mL).

### Genetic techniques

Oligonucleotide primers used in this study are listed in Table S1. PCR reactions for cloning and recombineering were performed with Q5 Hot Start High-Fidelity 2x Master Mix (NEB M0494S) according to manufacturer instructions. PCR products were purified using the Monarch® Spin PCR & DNA Cleanup Kit (NEB T1135) or gel extracted with the Monarch® Spin DNA Gel Extraction Kit (NEB T1120S) and sequenced by GENEWIZ at Azenta Life Sciences (South Plainfield, NJ). Plasmids were isolated using the Monarch Spin Plasmid Miniprep Kit (NEB T1110S) and sequence verified prior to transformation.

Plasmids generated for λ red recombineering (see below) and other chromosomal integration techniques were constructed through Gibson assembly (New England Biolabs E2621L) using traditional PCR or inverse (round-the-world) PCR using inward primers to introduce desired mutations or insertions following DpnI (NEB #R0176S) digestion. Constructed plasmids were electroporated into electrocompetent recipient strains or *E. coli* DH5α cells (ThermoFisher 11319019). Electrocompetent *Salmonella enterica* serovar Typhimurium were prepared by washing mid-log cultures with sterile cold water, mixing plasmid DNA with competent cells in 1 mm electroporation cuvettes, and electroporating at 1.8 kV. Transformed cells were recovered in 900 µL SOC medium for ≥2 hrs, pelleted & concentrated, and selected for on antibiotic-containing LB agar. Single colonies from selective plates were used for downstream validation and were saved for further experimentation.

### Phage enumeration and plaquing

Enumeration of phage particles via plaque isolation was performed using a double layer agar technique. ∼1 mL of bacterial culture containing phage was filtered through a 0.22 μm filter to remove bacterial cells and debris. The remaining lysate was diluted appropriately in sterile LB. Dilutions were applied in 5 μL drops to 3 mL 0.5% LB agar lawns seeded with 200 μL turbid indicator strain culture (∼10^8^ CFU). Inoculated plates were incubated overnight at 37°C.

### Determination of prophage viability & inducibility

Inducible and viable phages were detected in the supernatant of 4/74 prophage(+)/prophage(+_1_) after 24h culture. Where chemical induction was required, Mitomycin C (MitC, BPS Bioscience #27763) was added to newly inoculated cultures at a sub-inhibitory concentration (0.2 μg/mL). The double layer agar technique described above was used to test phage viability and enumeration using specific indicator strains to detect each phage. Inducibility, which ignores the ability of phages to reinfect new host cells, was tested by qPCR to detect successfully replicated prophages. Lysate samples from t=0 and t=3 hours were filtered (0.22 μm), treated with DNaseI at 37℃ for 10 minutes to eliminate remaining host DNA, and heated to 100℃ for 15 minutes to inactivate DNaseI and break open phage capsids.^51^ qPCR targeting unique prophage and prophage junction regions was performed using KAPA SYBR FAST qPCR Master Mix (2X) Universal (Roche KK4600) on a BioRad iCycler per manufacturer’s instructions with primers AZ11-14, 81-92. Unique prophage quantifications were normalized by junction values to normalize for remaining host genomic DNA. An increase in phage concentration between t=0 and t=3 implied successful phage induction.

### Construction of non-inducible prophages by pTOX allelic exchange

A prophage-induction knockout of the 4/74 prophage(+_1_) native phage Gifsy-2 was generated by allelic exchange targeting its induction antirepressor (*STM1019*). The pTOX-derived plasmid (pMS114), containing the deleted *STM1019* region, was electroporated into donor *E.coli* MFDλpir. Transformed cells were grown overnight in LB with 30 µM DAP + 2% glucose to repress pTOX toxin at 37℃ overnight. Recipient 4/74 prophage(+_1_) was incubated at 42℃ for 2 h to increase efficiency prior to pelleting.

Donor and recipient cells were pelleted and resuspended together in 10mM MgSO_4_ and washed twice with PBS. The final resuspension into 1xPBS + 30 µM DAP was plated entirely onto a 0.45 µm filter (MilliporeSigma Z290793) on LB with positive selection for the plasmid and incubated for 2-5 h. After conjugation, recipient cells were resuspended off the filter into a new tube containing 1 mL of MgSO_4_. These cells were pelleted, washed, and resuspended in 1 mL LB. Undiluted culture was plated onto LB + Cm34 + 2% glu. The rest of the culture was concentrated and plated onto the same selective media. Both plates were incubated at 37℃ for 24 h. Selected colonies were restreaked for purification on the same media and incubated at 37℃ overnight. Surviving cells were positive for the insertion of the pTOX construct into the genome. To perform reverse selection for the double crossover and removal of the pTOX2 backbone and the original genomic region, purified colonies were streaked onto 1-2% rhamnose LB plates and incubated for 12-14 h at 37℃. Remaining colonies were screened for the correctly substituted region via PCR (oligos AZ5-6) and further purified if needed. This generated the prophage-induction deficient strain 4/74 prophage(+_1_)_NI_.

### Construction of an *STM1019* complementation plasmid for *Salmonella* infection via Gibson assembly

To generate an induction complement, we used a plasmid construct to express the previously deleted region containing *STM1019* in our knockout background. We amplified the region deleted through pTOX allelic exchange (as described above) containing *STM1019*, its promoter, and its terminator as well as a pSC101 backbone using primers AZ16 & AZ17 to amplify the desired backbone region and AZ20 & AZ21 to amplify our target gene with homology into the target backbone. Regions were identified and confirmed using SalCom.^52^ Both amplified regions were combined by Gibson Assembly to generate the plasmid pAZ1. pAZ1 was electroporated into 4/74Δ prophage(+_1_)_NI_ to create the complemented knockout 4/74 prophage(+_1_)_NI_*^STM101^*^9^. Final strains were checked for correct assembly by sequencing prior to MIC testing.

### Antibiotic sensitivity testing & growth curves

Stocks of ciprofloxacin (CIP, Sigma-Aldrich #17850) suspended in water and 0.1M HCl, trimethoprim (TMP, Sigma Aldrich #T7883) suspended in DMSO, and MitC suspended in DMSO were sterilized using a 0.22 µm filter. For liquid culture susceptibility testing, antibiotics were added to OD_600_ = 0.01 bacterial cultures in LB at concentrations dependent on the experiment. To observe baseline strain growth rates, no antibiotic controls were done in replicate for each strain tested. Liquid assays were run in the BioTek Synergy H1 Multimode Reader (Agilent Technologies) or Sunrise^TM^ Absorbance Readers (TECAN #30190079) for 13-15 h at 37℃ with shaking, reading for OD_600_ every 10 minutes. Samples were normalized by blank LB only controls. Minimum inhibitory concentrations (MICs) were determined as the lowest concentration to yield final OD_600_ readings between OD_600_ = 0 and the minimum level of detection (OD_600_ ≈ 0.1). For each concentration, groups were compared using unpaired two-tailed multiple *t*-tests using the two-stage step-up Benjamini, Krieger, and Yekutieli procedure to control the false discovery rate (Q = 0.05), implemented in GraphPad Prism 10. For solid culture resistant mutant MIC tests, antibiotics were mixed with 2% LB agar at various concentrations prior to plate pouring. Agar mixtures were poured into OmniTrays (VWR International #62409) and dried in a sterile laminar flow hood. All cultures were inoculated using a 1.5 µL 96-well pin tool at full overnight concentration (OD_600_ ≥ 1.0). Plates were incubated at 37°C overnight, imaged, and analyzed manually. Pin tools were disinfected in 10% bleach, rinsed twice with water, and sterilized by 100% ethanol after each use. Relative MIC values on solid were determined by selecting the lowest concentration of CIP which inhibited colony growth and normalizing by the MIC value of their respective ancestral background. Relative and absolute MIC values were log₂-transformed prior to statistical analysis to account for MIC measurements obtained on a two-fold dilution scale. Group differences were assessed using Welch’s t-test.

### Construction of fluorescent *Salmonella* by λ Red recombineering

A construct containing a *recA* promoter-controlled GFP, an mScarlet-i3 controlled by a cuminic acid repressor (CymR)^53,54^, and 1000 base pairs of homology for each side of the target site was inserted at the Tn7 chromosomal integration site. The specified construct was generated via Gibson assembly (primers AZ55-60) and recombined into prophage(+), prophage(-), and prophage(+)/(-) derived CIP mutants containing the pSIM5-tet plasmid through λ Red recombination.^55,56^ The construct was inserted into the attTn7 region of each host with primers SP133 and SP134. The resulting strain was grown in LB supplemented with Amp/Carb100. Correct insertion at the attTn7 site was confirmed via PCR with primers AZ38 and AZ39.

### Fluctuation assays for baseline mutation rate

To determine the underlying mutation rate between ancestral strains, fluctuation tests were done as explained in the original Luria-Delbrück experiments^57^ and more recently by Rosche & Foster.^58^ Mutation rates and their corresponding error were then calculated using the FALCOR online tool.^59^

### Non-selective generation, quantification, and selection of antibiotic resistance mutants

5 clonal, biological replicates of 4/74 prophage(+) and 4/74 prophage(-) were inoculated into 5 mL of LB and incubated at 37℃ with shaking for 16 h. Incubation time was selected to generate mutants containing single/few resistance-conferring mutations (aiming to achieve “single-step evolution”).^60^ Antibiotic pressure was omitted during culture to prevent prophage induction and host death prior to mutation generation. All incubated cultures were challenged against a selective gradient of antibiotic on solid media (ranging from 0.01µg/mL to 0.08 ug/mL for CIP, 0.2 µg/mL to 1.8 µg/mL for MitC, and 0.2 µg/mL to 0.95 µg/mL for TMP). Counting plates were inoculated with diluted cultures for normalization. All plates were incubated for 48 h at 37℃ to allow for the appearance of slow growing mutants. Colony counts were collected using FIJI’s watershed segmentation analysis and particle annotation feature.^61^ Counts were normalized by dividing by the total number of cells plated for each respective replicate. Group differences of normalized values at each concentration were assessed using multiple unpaired Mann-Whitney U-tests with a two-stage step-up Benjamini, Krieger, and Yekutieli procedure to control the false discovery rate (Q = 0.05, implemented in GraphPad Prism 10). We selected and banked 9 CIP resistance mutants from one concentration plate (0.0625µg/mL CIP and 0.7µg/mL TMP) for each replicate in both the 4/74 prophage(+) and prophage(-) background. CIP selection concentration was chosen with consideration for the definition of “clinical resistance,” which is 0.06 µg/mL for *S.* Typhimurium.^24^ The TMP selection concentration was chosen to match the MIC fold change selected with CIP. A total of 143 samples for CIP and 38 samples for TMP were saved and used for further phenotypic and genotypic characterization.

### Whole genome sequencing & mutation identification

gDNA was extracted for each banked sample using the PureLink^TM^ Genomic DNA Mini Kit (Invitrogen^TM^ #K182002). All samples were normalized to 0.5 ng/µL and used for sequencing library preparation as described in Baym, et al.^62^ Some samples were library prepped a second time for resequencing using the Twist Enzymatic Fragmentation kit (Twist Biosciences # 104206). All samples were multiplexed using custom designed 8 base pair primers (IDT) to generate paired-end reads. All quantification steps during library prep were done using the Quant-It^TM^ dsDNA Assay Kit (broad-range, Invitrogen^TM^ #Q33267). Pooled and final libraries were quantified by qPCR with the KAPA Library Quantification Kit (Roche #4844) on the BioRad^®^ iCycler. Completed pooled library samples were sent to the Bauer Core Facility at Harvard University for sequencing. Mutations were identified by aligning the generated short read sequences to an ancestral reference (GenBank CP002487) using *breseq*.^63^ As these mutations are clonal, mutations with an observed frequency of ≥ 85% and coverage over 20x were kept for mutant identification (Fig. S4). Single mutations were immediately characterized as resistance-conferring. Others were classified as putatively resistance-conferring if they were recurrent across independent isolates, previously characterized as resistance-associated, or occurred in genes with established roles in resistance to the selecting antibiotic. Synonymous mutations and variants lacking recurrence or functional support were ignored or categorized as putative compensatory mutations. Putative resistance-conferring mutations were grouped into pathway categories according to the known function of the affected gene, with genes participating in shared resistance mechanisms or cellular processes assigned to the same class. Pathway assignments were based on published functional annotations and literature curation. Implication in clinical resistance was assigned to mutations which appeared in the Comprehensive Antibiotic Resistance Database^26^ or were observed clinically in literature.^5,25,29,50,64,65^ Differences in pathway distributions between backgrounds were assessed using a permutation-based chi-square test with 10,000 random label shuffles, recalculating the chi-square statistic at each iteration to generate an empirical null distribution. Pathway enrichment was tested using Fisher’s exact tests and a Monte Carlo permutation framework in which group labels were randomly reassigned 10,000 times to generate empirical null distributions.

### Flow cytometry of fluorescent SOS strains

21 clonal cultures (3 independent, biological replicates) of CIP resistant mutants and ancestral strains (4/74 prophage(+) & 4/74 prophage(-)) were transformed with attTn7::precA-GFPova, mScarlet-i3-cymR, ampR; were inoculated into 1.5 mL LB + Amp100 in a 96-deep well plate and incubated at 37℃ with shaking overnight. Overnights were also generated for several flow control samples (non-fluorescent, mScarlet-i3-cymR only, and precA-GFPova only). 3 hours prior to flow cytometry, cultures were split into 2 plates: +/- CIP. Cultures were back diluted 1:100 into a 96 well plate containing fresh LB medium and 100 uM cym (200 uL total, omitting the non-fluorescent and GFPova controls). +CIP plates had 0.06 ug/mL CIP added to each well for resistant mutants and 0.01 ug/mL CIP for ancestral and GFPova only wells (omitting the non-fluorescent and mScarlet-i3 controls). These concentrations are both sub-MIC for their respective cultures. After a 3 h, 37℃ incubation, cultures were diluted 1:100 into 1xPBS in a 96 well plate (150 uL total). Flow cytometry was performed on the BD FACSymphony™ A1 Cell Analyzer (BD BioSciences) using HTS settings. Data collection was done using BD FACSDiva software (BD BioSciences) and analyzed using FlowJo v10.10 (BD). Differences in fold change GFP expression (both normalized and raw) were assessed using two-tailed Mann–Whitney U tests implemented in Python.

### Fluorescent antibiotic drop tests

Observation of SOS response induction from various mechanism antibiotics was performed through antibiotic drop tests. On 25 mL LB + 2% agar plates, 5 µL of working stock antibiotics were pipetted into the middle of their respective agar plates. Antibiotics were allowed to absorb completely next to a flame. Plates were left upright at 4℃ for 5-18 hours to allow for full antibiotic diffusion. Overnight cultures were prepared from clonal samples of our fluorescent SOS response strains of 4/74 prophage(+) and 4/74 prophage(-) in LB + Amp100. These were left shaking at 37℃ until turbid (>OD 1.0). 100 µL of overnight culture was plated at full concentration onto each fully diffused antibiotic plate and spread using sterile glass beads. These plates were dried and incubated at 37℃. Plates were imaged the next day using the custom fluorescent imager as described in Rand, et al. under both brightfield and GFP settings.^66^ SOS induction was characterized by the expression of GFP at the zone of inhibition and the surrounding bacterial lawn.^45^

## Acknowledgements

We are grateful to past and present members of the Baym Laboratory at Harvard Medical School for helpful discussions and materials. We thank Nicholas Wenner, Molly Sargen for several bacterial strains used in this study. We thank the Bauer Core at Harvard University for sequencing support and expertise throughout this project. We are grateful to the Laboratory Operations team in the Harvard Program in Therapeutic Sciences (HiTS) for essential operational support, in particular Tenzin Phulchung, Nicole Anderson and Scott Slimmer. We additionally thank Twist Biosciences for their generous sequencing support through the Academic Core Grant Program. This work was supported by the NIGMS of the National Institutes of Health (R35GM133700 and R35GM156320), the David and Lucile Packard Foundation, the Pew Charitable Trusts, and NSF grant IOS-2331228. A.D.Z. acknowledges support from the Systems, Synthetic, and Quantitative Biology PhD program training award (T32GM135014) and the Graduate Prize Fellowship.

