## Supplementary Tables and Figures for "SOS-mediated prophage induction constrains resistance evolution to DNA-damaging antibiotics"

**Supplementary Table 1 (Table S1). Oligonucleotide primers used in this study.**
Overhangs are in lowercase text letters.

| Oligo ID | Name | Sequence (5’ → 3’) | Purpose |
|---|---|---|---|
| AZ5 | gifsy2NI_fwd | CACACTAGCCTTTGCAAAAAG | Amplification of 734bp region to confirm pTOX allelic exchange deleting *STM1019* |
| AZ6 | gifsy2NI_rev | TGGCGCATAATTGTCAGGTTA | |
| AZ11 | gifsy2_fwd | GAAGTGGCTGAGAGGG | Amplification of 286bp region of gifsy-2 prophage |
| AZ12 | gifsy2_rev | GCTATCATTCCTCCTGCTGA | |
| AZ16 | pSC101_Km_fwd | ACCTGCAGGCATGCAA | Amplification of 4.2kb backbone of pSC101 for Gibson assembly |
| AZ17 | pSC101_Km_rev | GACGTGGTGTTAGCTGTG | |
| AZ18 | STM1019_fwd | TCAAGGATACCAGAGGAGATGT | Amplification of 1170 bp region capturing the DinI-like homolog in 4/74 |
| AZ19 | STM1019_rev | TGACTGCAGAATTTCGCTG | |
| AZ20 | STM1019_pSC101_fwd | ACCACAGCTAACACCACGTCTCAAGGATACCAGAGGAGATGT | Amplification of 1170 bp region including STM1019 in 4/74; overhang into pSC101 for Gibson assembly |
| AZ21 | STM1019_pSC101_rev | GAGCTTGCATGCCTGCAGGTtgactgcagaatttcgctg | |
| AZ22 | rbs+mscar_fwd | tttaagaaggagatatacatATGGACTCAACCGAAGC | Amplification of the 730 bp mScarlet-i3 gene block; overhang containing an RBS |
| AZ23 | mscar_rev | TTAGCTGCCTCCGCTT | |
| AZ24 | rbs_cym_fwd | atgtatatctccttcttaaaTACAAACAGACCAGATTGTCTGTTT | Amplification of 4460 bp PCV3 backbone; overhang into a rbs+mscar construct for pAZ3 construction via Gibson assembly |
| AZ25 | mscar_cym_rev | aagcggaggcagctaaCCTGCAGGCATGCAAG | |
| AZ31 | pcym_rev | GTCCCTATCTGCTGCCCTAG | Amplification of 1759 bp region in pAZ3 containing cymR & mscar motifs; overhang into pFCiGrecA for pAZ4 construction via Gibson assembly |
| AZ34 | cymterm_ampR_fwd_redo | tctcatgaccaaaatcccttaacgtgTGCATAATGTGCCTGTCAAATGGA | |
| AZ38 | Sal474_Tn7_external_F | CGATCCACCTGGATTGAGCC | Amplification of the Tn7 region to check for recombination in 4/74 |
| AZ39 | Sal474_Tn7_external_R | CTTCTACACCGTTCCGCTGC | |
| AZ35 | pfcigreca3_pcym_ | ctagggcagcagatagggacCATTAAA | Amplification of 3375 bp |

| | | | |
|---|---|---|---|
| | fwd | GCTTTCGCGGACTGC | region of pFCiGrecA containing precA-GFPova fusion, AmpR, & ORI; overhang into pAZ3 for pAZ4 construction via Gibson assembly |
| AZ40 | pfcigrecaORI_rev | GGGCTGTGAGCGCTCTTC | |
| AZ44 | 10kbhomol_ori_R | TGAGTTTTCGTTCCACTGAGC | Amplification of 845 bp region of pAZ4 containing the ORI for pAZ5 construction |
| AZ45 | 10kbhomol_ori_F | TTATGCAGGGCTGTGAGCG | |
| AZ57 | Tn7right_pAZ4ampR_R | tttttgataatctcatgaccaaaatcccttaacgCTTCTGCCTGGTACTACATTGTAC | Amplification of 1077 bp Tn7 right arm from 4/74; overhangs into both the ORI & AmpR of pAZ4 for Gibson assembly of pAZ5 |
| AZ58 | Tn7rightStart_pAZ4ori_F | cttttctacggggtctgacgctcagtggaacgaaaactcaCACGCCGACGAACTGCTGTC | |
| AZ59 | Tn7leftend_pAZ4cym_F | tttgcagaatccctgcttcgtccatttgacaggcacaGTCGACAGACGGCCTTTTTTGT | Amplification of 1041 bp Tn7 left arm from 4/74; overhangs into both the ORI & cymR of pAZ4 for Gibson assembly of pAZ5 |
| AZ60-2 | Tn7leftStart_pAZ4ori_R | agtcagtgagcgaggaagcggaagagcgctcacagCTGCTGGATGAAGTGGAGGTC | |
| SP133 | Sal474_Tn7_10kbhomol_recombineer_F | CTGCTGGATGAAGTGGAGGTC | Amplification of 6285 bp region of pAZ5 with 1000 bp of homology to 4/74 Tn7 region for lambda red recombination into 4/74 |
| SP134 | Sal474_Tn7_10kbhomol_recombineer_R | CGCCGACGAACTGCTGTC | |

**Supplementary Table 2 (Table S2). Unique mutations of single-step evolved 4/74 prophage(+) & prophage(-)**

*indicates mutated genes only seen in tandem with other mutations

| Mutated gene(s) | Mutant background(s) | Mutation(s) | Mutation location (CP002487) |
|---|---|---|---|
| **DNA gyrase & topoisomerase** | | | |
| *gyrA* | 4/74 prophage(+)<br>4/74 prophage(-) | T → C<br>D87G | 2,373,804 |
| | 4/74 prophage(-) | C → A<br>D87Y | 2,373,805 |
| | 4/74 prophage(+) | G → A<br>S83F | 2,373,816 |
| | 4/74 prophage(-) | G → T<br>A119E | 2,373,708 |
| | 4/74 prophage(-) | C → T<br>D82N | 2,373,820 |
| | 4/74 prophage(+) | Δ12bp<br>in frame | 2,372,225 |
| *gyrB* | 4/74 prophage(+) | T → G<br>E466D | 4,061,187 |
| | 4/74 prophage(+) | T → A<br>E466D | 4,061,187 |
| | 4/74 prophage(+) | G → A<br>S464F | 4,061,194 |
| **Efflux pumps & membrane proteins** | | | |
| *soxR* | 4/74 prophage(+) | G → C<br>D137H | 4,526,081 |
| | 4/74 prophage(+) | G → A<br>G145D | 4,526,106 |
| | 4/74 prophage(+)<br>4/74 prophage(-) | T → C<br>L149P | 4,526,118 |
| | 4/74 prophage(+) | C → T<br>R20C | 4,525,730 |
| | 4/74 prophage(+) | Δ27bp | 4,526,082 |

| | | | |
|---|---|---|---|
| | | in frame | |
| | 4/74 prophage(-) | A → G<br>D137G | 4,526,082 |
| | 4/74 prophage(-) | G → T<br>D137Y | 4,526,081 |
| *ramR* | 4/74 prophage(+) | C → A<br>G42V | 638,091 |
| | 4/74 prophage(+) | A → C<br>L54R | 638,055 |
| *envZ* | 4/74 prophage(+) | C → A<br>P248Q | 3,680,389 |
| | 4/74 prophage(+) | G → T<br>P248Q | 3,680,389 |
| *ftsI2** | 4/74 prophage(+)<br>4/74 prophage(-) | C → A<br>K3N | 1,892,887 |
| *sapB** | 4/74 prophage(-) | A → G<br>T137A | 1,742,586 |
| **Transcription/ Replication** | | | |
| *rpoA* | 4/74 prophage(-) | C → T<br>C269Y | 3,604,967 |
| *rpoC* | 4/74 prophage(+)<br>4/74 prophage(-) | G → A<br>G1354E | 4,395,405 |
| | 4/74 prophage(+) | G → A<br>G1354R | 4,395,404 |
| *rpoD* | 4/74 prophage(+) | Δ30bp<br>T526M | 3,398,547 |
| | 4/74 prophage(+) | A → G<br>N430S | 3,398,259 |
| *deaD* | 4/74 prophage(+) | Δ13bp<br>frameshift | 3,468,025 |
| *rcsA** | 4/74 prophage(+) | G → C<br>R47C | 2,025,135 |
| *ygfE** | 4/74 prophage(-) | A → C<br>F191V | 2,607,731 |
| **Stress response** | | | |

| | | | |
|---|---|---|---|
| *clpP* | 4/74 prophage(+) | C → T<br>Q87* | 502,837 |
| | 4/74 prophage(+) | Δ5bp<br>frameshift | 503,018 |
| **Metabolism** | | | |
| *purB* | 4/74 prophage(+) | G → T<br>L337M | 1,276,702 |
| | 4/74 prophage(-) | $(ATAAGCG)_{2\rightarrow1}$<br>frameshift | 1,276,647 |
| | 4/74 prophage(-) | $(ATAAGCG)_{2\rightarrow3}$<br>frameshift | 1,276,654 |
| | 4/74 prophage(-) | +GATAAGC<br>frameshift | 1,276,638 |
| | 4/74 prophage(-) | +CGCCACGTT<br>in frame | 1,277,146 |
| | 4/74 prophage(-) | C → T<br>W43* | 1,277,582 |
| | 4/74 prophage(-) | T → G<br>Q390P | 1,276,542 |
| | 4/74 prophage(-) | T → G<br>T170P | 1,277,203 |
| | 4/74 prophage(-) | +A<br>frameshift | 1,276,511:1 |
| | 4/74 prophage(-) | G → A<br>Q38* | 1,277,599 |
| | 4/74 prophage(-) | +G<br>frameshift | 1,277,682:1 |
| | 4/74 prophage(-) | T → G<br>T409P | 1,276,486 |
| | 4/74 prophage(-) | Δ10bp<br>frameshift | 1,276,635 |
| *ribD* | 4/74 prophage(+) | Δ1bp<br>frameshift | 469,772 |
| *ribH* | 4/74 prophage(-) | T → C<br>L151S | 470,973 |
| *icdA* | 4/74 prophage(+) | Δ54 bp<br>in frame | 1,282,084 |
| *eutC** | 4/74 prophage(+) | Δ1bp | 2,564,148 |

| | | | |
|---|---|---|---|
| | | frameshift | |
| *ackA** | 4/74 prophage(+) | A → C<br>T233P | 2,446,351 |
| *aceF** | 4/74 prophage(+) | Δ1bp<br>S394A<br>frameshift | 180,101 |
| *trpD** | 4/74 prophage(-) | G → T<br>G356* | 1,778,030 |
| *lipB** | 4/74 prophage(-) | Δ1bp<br>frameshift | 696,930 |
| | 4/74 prophage(-) | +AG<br>frameshift | 696, 933 |
| **Motility** | | | |
| *trmE** | 4/74 prophage(-) | Δ13bp<br>frameshift | 4,070,004 |
| **Large-scale deletions** | | | |
| *[mnma], ymfB, STM474_1234, ymfC, icdA, cspH, [envF]* | 4/74 prophage(+) | Δ6,773bp | 1,279,043 |
| *[purB], hfld, mnma, ymfB, STM474_1234, ymfC, icdA, STM474_1237, envF, msgA, envE, cspH, pagD, pagC, STM474_t1245, [pliC]* | 4/74 prophage(+) | Δ14,353bp | 1,276,676 |
| *[galU], hnr, [rssa]* | 4/74 prophage(+) | Δ2,847bp | 1,804,872 |
| **Intergenic** | | | |
| *yiaG → / → cspA* | 4/74 prophage(+) | T → A<br>(+202/-87) | 3,857,668 |
| | 4/74 prophage(+) | A → G<br>(+176/-113) | 3,857,642 |
| *STM474_2610 ←/← guaA** | 4/74 prophage(+) | G → T<br>(-106/+51) | 2,260,470 |
| *sulA ←/→ yccr** | 4/74 prophage(-) | +C<br>(-45/-172) | 1,119,250:1 |

**Supplementary Table 3 (Table S3). Ancestral & selected mutant identities**
(more detailed information about mutations is available in Supplementary Table 2)

| Mutant/sample number | Strain background & replicate | Mutated/affected gene(s) | Mutation | Sequencing coverage | CIP MIC (µg/mL) |
|---|---|---|---|---|---|
| 4/74 prophage(+) | prophage(+) | n/a | n/a | n/a | 0.025 |
| s2 | prophage(+) replicate 3 | *[purB], hfld, mnma, ymfB, STM474_1234, ymfC, icdA, STM474_1237, envF, msgA, envE, cspH, pagD, pagC, STM474_t1245, [pliC]* | Δ14,353 bp: 1,276,676 | 34.3x | 0.0975 |
| s11 | prophage(+) replicate 1 | *gyrA* | G → A S83F | 264x | 0.44 |
| s12 | prophage(+) replicate 1 | *gyrA* | G → A S83F | 232x | 0.44 |
| s13 | prophage(+) replicate 4 | *gyrA* | G → A S83F | 69x | 0.88 |
| s14 | prophage(+) replicate 4 | *envZ* | C → A P248Q | 268x | 0.0625 |
| s15 | prophage(+) replicate 4 | *icdA; deaD* | Δ54bp: 1,282,084; Δ13bp: 3,468,025 | 267.9x | 0.0625 |
| s16 | prophage(+) replicate 4 | *ramR; ftsI2* | C → A G42V; C → A K3N | 73.5x | 0.08 |
| s17 | prophage(+) replicate 5 | *rpoC; eutC* | G → A G1354R | 38.3x | 0.22 |
| s18 | prophage(+) replicate 5 | *gyrA* | T → C D87G | 46x | 0.44 |
| s19 | prophage(+) replicate 5 | *gyrA* | T → C D87G | 20x | 0.44 |
| s99 | prophage(+) replicate 3 | *gyrA* | T → C D87G | 55x | 0.22 |
| s102 | prophage(+) replicate 2 | *clpP* | Δ5bp: 503,018 | 74.8x | 0.0625 |
| s103 | prophage(+) replicate 4 | *soxR* | G → C D137H | 40.4x | 0.22 |
| s106 | prophage(+) replicate 1 | *rpoD; ftsI2* | Δ30bp: 3,398,547; C → A | 88x | 0.0975 |

| | | | | | |
|---|---|---|---|---|---|
| | | | K3N | | |
| s107 | prophage(+) replicate 3 | *gyrB* | T → A E466D | 21x | 0.135 |
| s110 | prophage(+) replicate 2 | *soxR* | T → C L149P | 20x | 0.08 |
| s111 | prophage(+) replicate 4 | *clpP* | C → T Q87* | 73.5x | 0.08 |
| s114 | prophage(+) replicate 1 | *gyrB* | T → G E466D | 262X | 0.135 |
| s115 | prophage(+) replicate 3 | *[galU], hnr, [rssa]* | Δ2,847 bp: 1,804,872 | 74.5x | 0.0975 |
| s118 | prophage(+) replicate 2 | *rpoC* | G → A G1366E | 67.1x | 0.08 |
| s121 | prophage(+) replicate 1 | *gyrB* | G → A S464F | 406.3x | 0.22 |
| s122 | prophage(+) replicate 4 | *ribD* | Δ1bp | 174x | 0.115 |
| s128 | prophage(+) replicate 1 | *yiaG → / → cspA* | T → A (+202/-87) | 200x | 0.135 |
| s129 | prophage(+) replicate 4 | *yiaG → / → cspA* | A → G (+176/-113) | 304x | 0.22 |
| s131 | prophage(+) replicate 1 | *soxR; purB* | Δ27bp: 4,526,082; G → T L377M | 296x | 0.0975 |
| s132 | prophage(+) replicate 1 | *gyrA* | T → C D87G | 63.8x | 0.22 |
| s142 | prophage(+) replicate 4 | *[mnma], ymfB, STM474_1234, ymfC, icdA, cspH, [envF]* | Δ6,773 bp: 1,279,043 | 25x | n.d. |
| s147 | prophage(+) replicate 5 | *soxR; rcsA* | C → T Q19H; G → C Q47H | 34.3x | 0.135 |
| s149 | prophage(+) replicate 1 | *rpoD* | A → G N430S | 452x | 0.22 |
| s150 | prophage(+) replicate 4 | *gyrA; soxR; ackA* | Δ12bp: 2,372,225; G→A G145D; A → C T233P | 304x | 0.22 |
| s153 | prophage(+) replicate 5 | *soxR; rcsA; aceF* | C → T R20C; G → C Q47H; | 191.9x | 0.135 |

| | | | | | |
|---|---|---|---|---|---|
| | | | Δ1bp: 180,101 | | |
| s154 | prophage(+)<br>replicate 4 | *ramR; envZ;*<br>*STM474_2610*<br>←/← *guaA* | A → C<br>L54R;<br>G→ T<br>P248Q;<br>G → T<br>(-106/+51) | 368.8x | 0.0625 |
| 4/74<br>prophage(-) | prophage(-) | Δ*Gifsy-2*<br>Δ*ST64B*<br>Δ*Gifsy-1*<br>Δ*SopE*<br>*minE*→/← *rnd*<br><br>*iolC* | Δ45,596bp: 1,054,761<br>Δ40,147: 2,039,803<br>Δ51,187bp: 2,726,265<br>Δ45,069bp: 2,855,501<br>G→A<br>(+108/+14)<br>(G)$_{8\rightarrow 7}$ | n/a | 0.035 |
| s3 | prophage(-)<br>replicate 1 | *gyrA* | G → A<br>D82N | 147.9x | 0.22 |
| s4 | prophage(-)<br>replicate 4 | *purB* | T → G<br>T170P | 274x | 0.0975 |
| s20 | prophage(-)<br>replicate 1 | *gyrA* | C → A<br>D87Y | 27.4x | 0.22 |
| s21 | prophage(-)<br>replicate 1 | *purB; ftsI2* | (ATAAGCG)$_{2\rightarrow 1}$;<br>C → A<br>K3N | 97.9x | 0.0975 |
| s22 | prophage(-)<br>replicate 1 | *purB; ftsI2* | +GATAAGC;<br>C → A<br>K3N | 44.8x | 0.0975 |
| s25 | prophage(-)<br>replicate 1 | *gyrA* | C → T<br>D82N | 28.7x | 0.22 |
| s26 | prophage(-)<br>replicate 1 | *rpoC* | G → A<br>G1354Q | 57.9x | 0.135 |
| s27 | prophage(-)<br>replicate 1 | *purB; ftsI2* | (ATAAGCG)$_{2\rightarrow 1}$;<br>C → A<br>K3N | 80.6x | 0.0975 |
| s28 | prophage(-)<br>replicate 2 | *purB; sapB;*<br>*sulA* ←/→ *yccR* | T → G<br>T409P;<br>A → G<br>T137A;<br>+C (-45/-172) | 380x | 0.0975 |
| s30 | prophage(-)<br>replicate 2 | *purB* | +GATAAGC | 71.2x | 0.0975 |
| s31 | prophage(-)<br>replicate 2 | *purB; trpD* | G → A<br>Q38*<br>G → T<br>G356* | 20x | 0.0975 |

| | | | | | |
|---|---|---|---|---|---|
| s32 | prophage(-) replicate 2 | *purB* | C → T W43* | 60.3x | 0.0975 |
| s33 | prophage(-) replicate 2 | *purB* | +GATAAGC | 23.9x | 0.0975 |
| s34 | prophage(-) replicate 2 | *purB* | T → G T409P | 299x | 0.0975 |
| s35 | prophage(-) replicate 2 | *purB; ftsI2* | T → G Q390P; C → A K3N | 37.9x | 0.0975 |
| s36 | prophage(-) replicate 3 | *gyrA* | G → T A119E | 37.2x | 0.22 |
| s37 | prophage(-) replicate 3 | *purB* | +GATAAGC | 49.2x | 0.115 |
| s38 | prophage(-) replicate 3 | *purB* | +GATAAGC | 53.4x | 0.0975 |
| s40 | prophage(-) replicate 3 | *gyrA* | T → C D87G | 57.2x | 0.22 |
| s41 | prophage(-) replicate 3 | *purB; ribH; trmE* | Δ10bp: 1,276,365; T → C L151S; Δ13bp: 4,070,004 | 218x | 0.08 |
| s42 | prophage(-) replicate 3 | *soxR* | T → C L149P | 43.6x | 0.22 |
| s43 | prophage(-) replicate 4 | *soxR* | G → T D137Y | 47.5x | 0.135 |
| s44 | prophage(-) replicate 4 | *purB* | +G | 315x | 0.0625 |
| s45 | prophage(-) replicate 4 | *soxR; yfgE* | G → T D137Y; A → C F191V | 165x | 0.22 |
| s47 | prophage(-) replicate 4 | *soxR* | A → G D137G | 351x | 0.135 |
| s48 | prophage(-) replicate 4 | *soxR* | G → T D137Y | 51.2x | 0.22 |
| s49 | prophage(-) replicate 4 | *purB* | +A | 91.8x | 0.08 |
| s50 | prophage(-) replicate 5 | *purB; ftsI2* | +GATAAGC; C → A K3N | 71.5x | 0.115 |
| s51 | prophage(-) replicate 5 | *purB* | +GATAAGC | 93.6x | 0.08 |

| | | | | | |
|---|---|---|---|---|---|
| s140 | prophage(-) replicate 3 | *soxR* | T → C L149P | 56.7x | n.d. |
| s143 | prophage(-) replicate 5 | *rpoA* | C → T C269Y | 25x | 0.135 |
| s144 | prophage(-) replicate 5 | *purB* | +CGCCACGTT | 266x | 0.08 |
| s156 | prophage(-) replicate 5 | *purB; lipP* | +ATAAGCG; Δ1bp: 696,930, +AG | 89.4x | 0.44 |

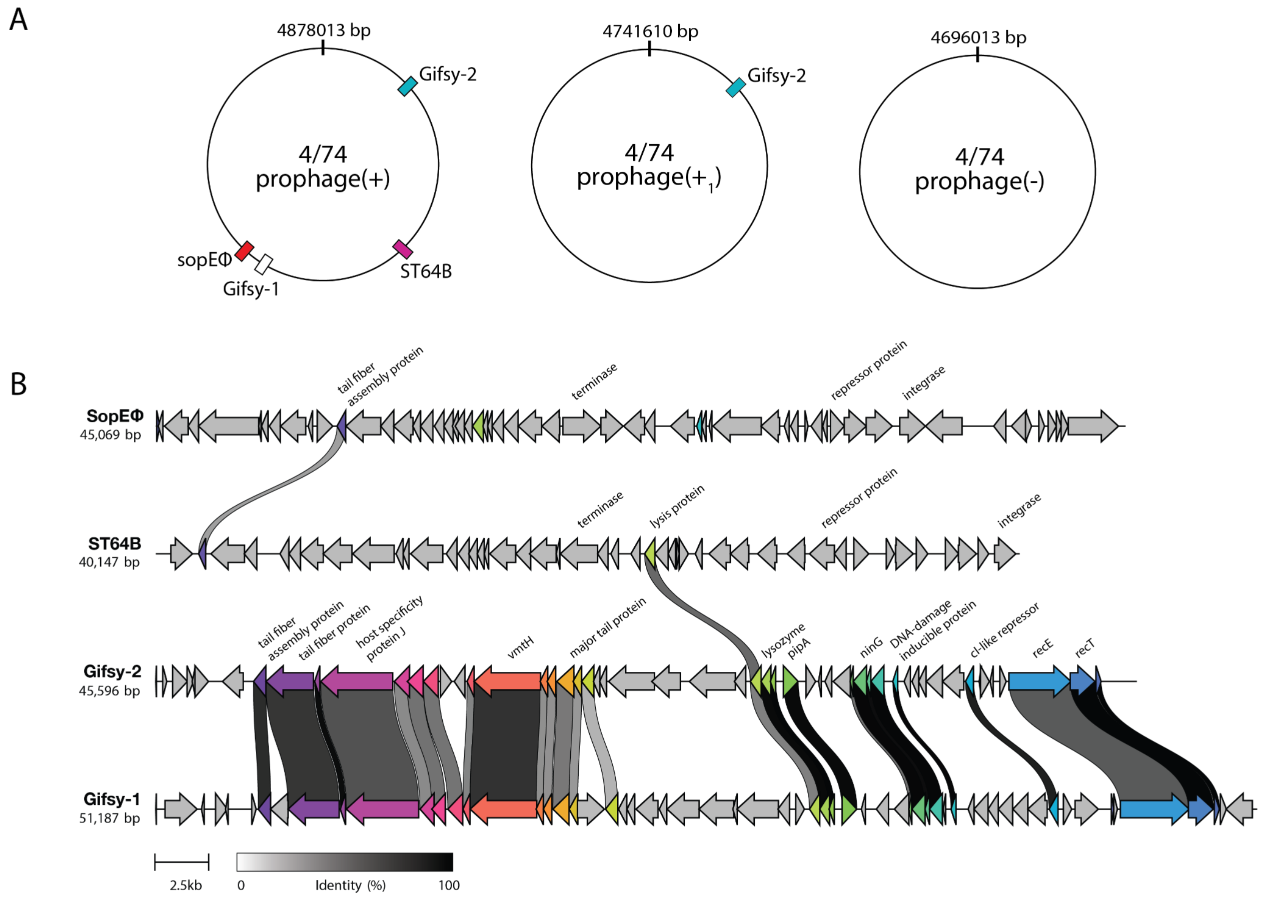


**Supplementary Figure 1 (S1).** ***Salmonella*** **Typhimurium 4/74 prophage composition & synteny.**

Prophage regions were identified within each variant of 4/74 and were internally compared to assess similarity. **A.** Schematic representation of prophage composition across *S.* Typhimurium variants used in this study, highlighting the location of each mobile genetic element. **B.** Comparative synteny analysis generated using clinker, showing homologous protein clusters and gene similarity across each integrated prophage. Genes are represented as directional arrows with homologous genes grouped by shared sequence similarity and colored accordingly.

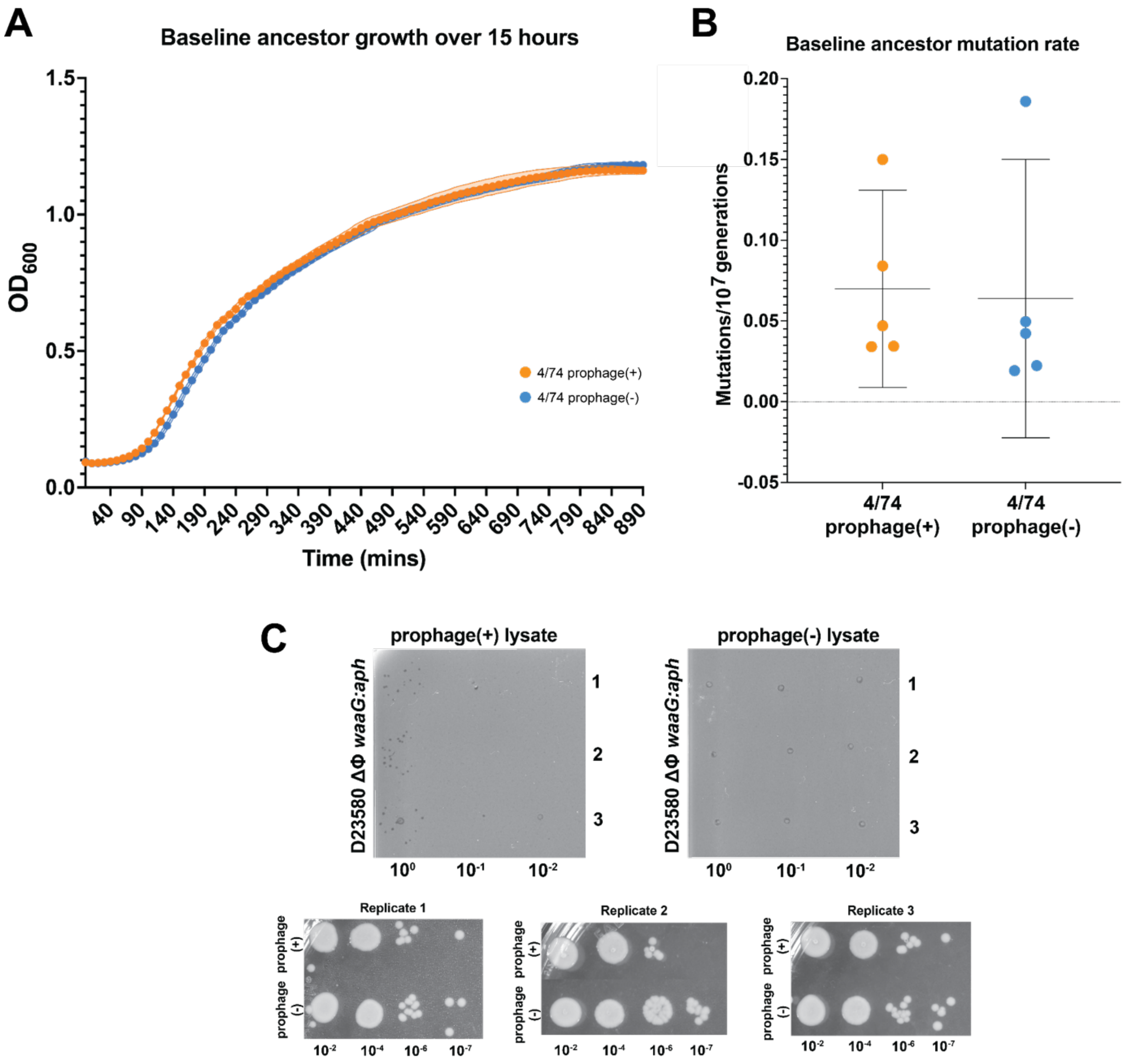


**Supplementary Figure 2 (S2). There are no prophage-dependent differences across growth and spontaneous mutation rate for ancestral *Salmonella* Typhimurium.**

**A.** Growth curves of lysogenic 4/74 prophage(+) and isogenic 4/74 prophage(-). Colored points represent OD600 measurements collected every 10 min for 13 h across four biological replicates. No measurable differences in growth rate were observed between backgrounds. **B.** Fluctuation assay calculations of spontaneous mutation rates per $10^{7}$ bacterial generations. Each point represents the estimated mutation rate (see Methods) for an individual biological replicate colored by ancestral background. There was no measurable difference in mutation rate between strains. **C.** Representative plaque assays and corresponding colony counts used to quantify spontaneous prophage induction following 24 h incubation. Filtered lysates were spotted onto lawns of indicator strain *Salmonella enterica* serovar Typhimurium strain D23580ΔΦ *waaG:aph*. Plaque- and colony-forming units were used to calculate a spontaneous induction rate of $6 \times 10^{-6}$ PFU/CFU for 4/74 prophage(+).

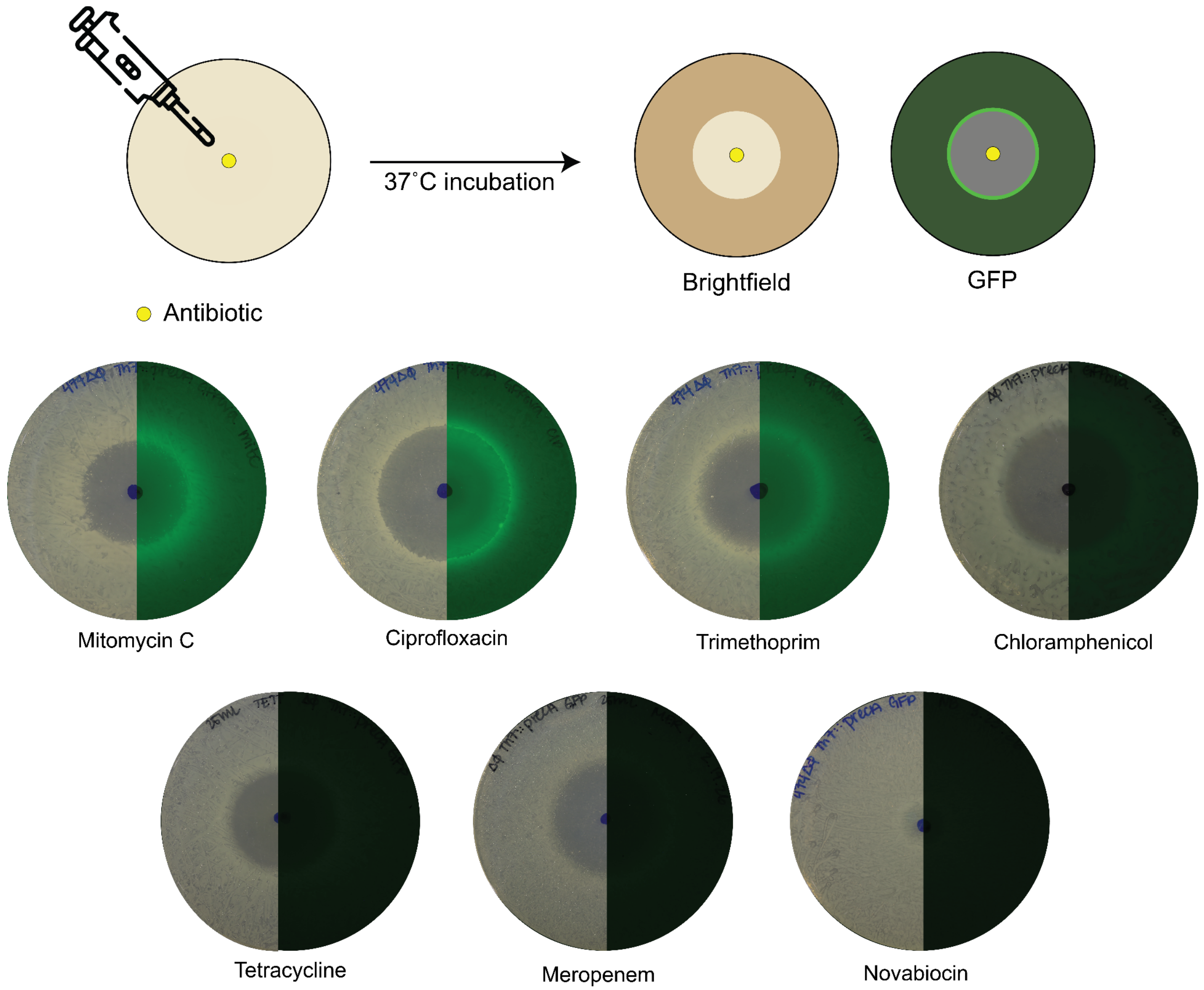


**Supplementary Figure 3 (S3). SOS response induction is dependent on antibiotic class and mechanism of action.**

Drop assays demonstrating SOS response activation through *precA* reporter expression. Concentrated antibiotics (5 µL) were applied to the center of 25 mL 2% LB agar plates and allowed to diffuse for >5 h prior to inoculation with bacterial lawns carrying a chromosomal *precA*-GFPova reporter. Plate images are shown as half brightfield and half GFP fluorescence. SOS activation is indicated by GFP fluorescence surrounding and extending beyond the zone of inhibition.

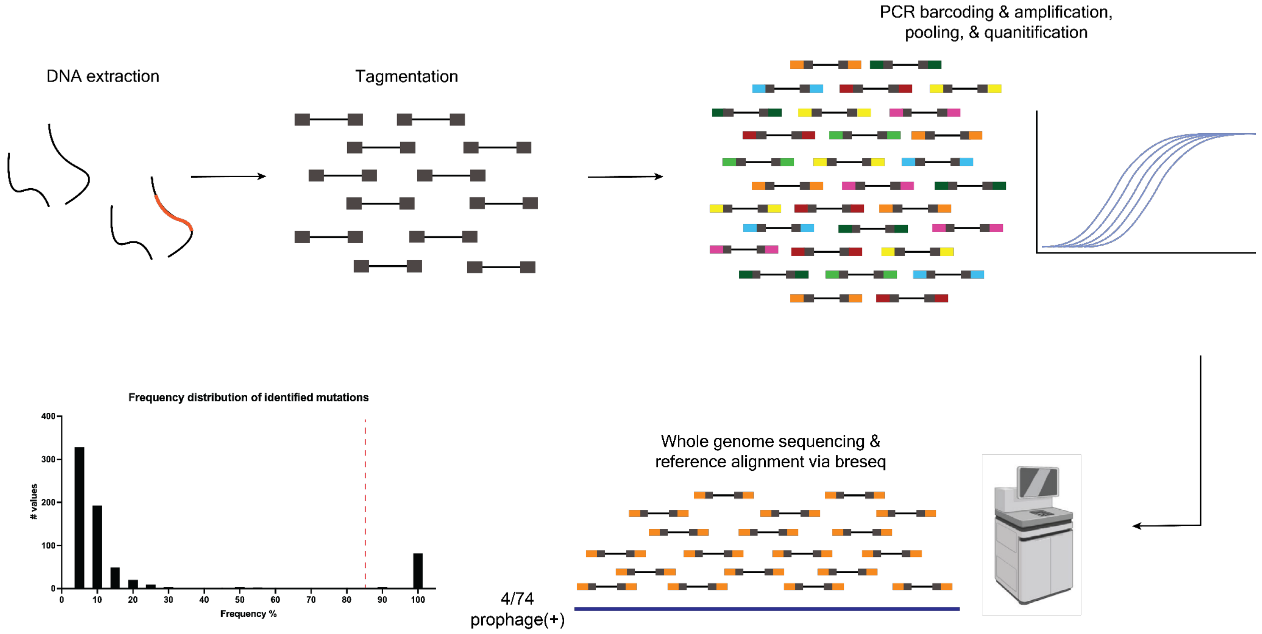


**Supplementary Figure 4 (S4). Whole genome sequencing alignment via *breseq* provides high coverage mapping to reference genomes**

Workflow for whole-genome sequencing and analysis of CIP-resistant isolates. Genomic DNA was extracted from clonal overnight cultures and tagmented as described by Baym et al. Unique dual 8 bp barcodes were added to tagged DNA fragments by PCR using custom primers (Supplementary Table 1). Amplicons were pooled, quantified by qPCR, and sequenced at the Harvard Bauer Core Facility. Sequencing reads were quality controlled using FastQC, preprocessed with Trimmomatic, aligned using minimap2 and SAMtools, and analyzed with *breseq* against the 4/74 WT reference genome. Because isolates were clonal, mutation calling thresholds were set at 85% allele frequency.

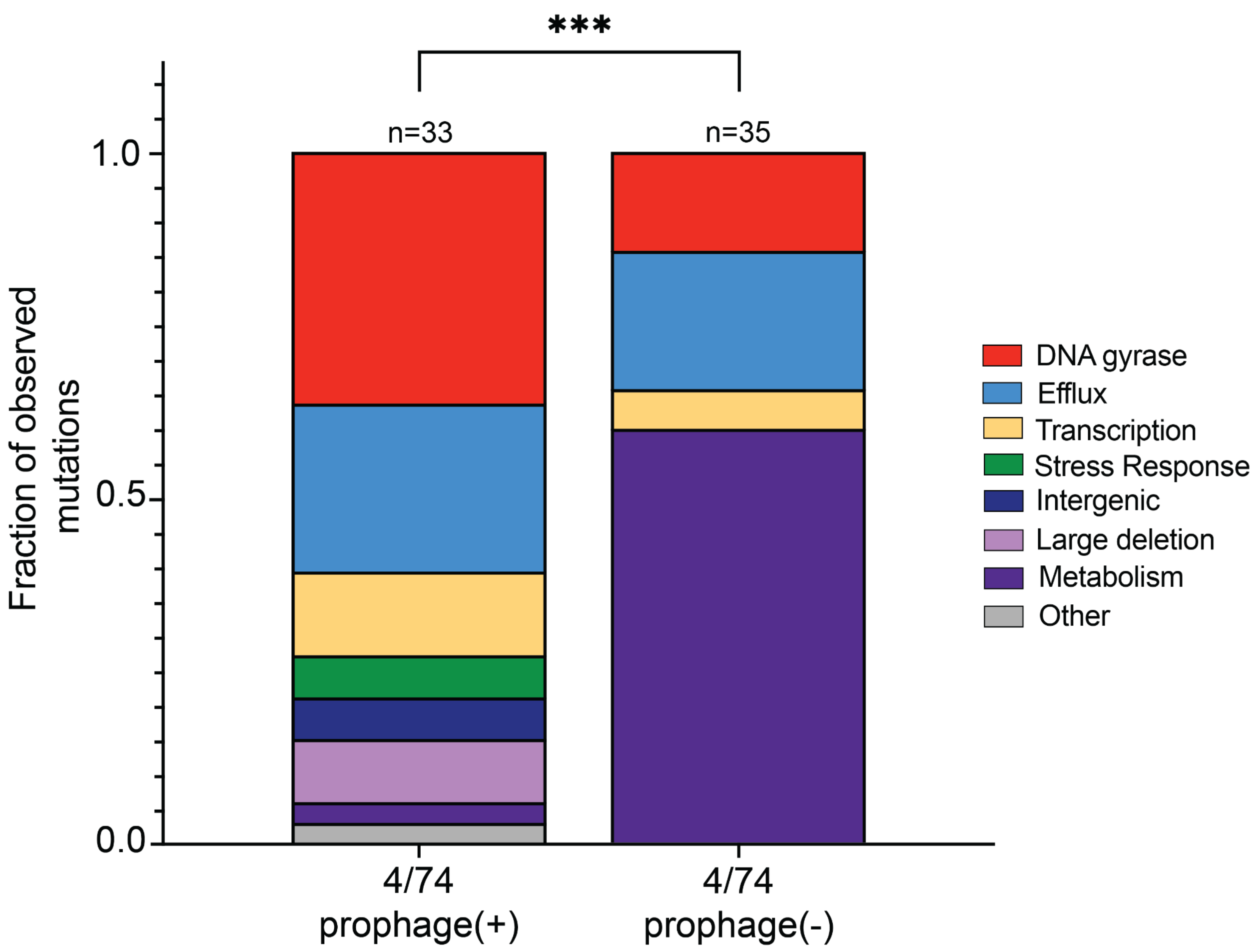


**Supplementary Figure 5 (S5). Absolute mutated pathway distributions also differ by prophage carriage.**

Bar plots showing the distribution of overall observed pathways/categories represented by genes carrying resistance-conferring mutations in 4/74 prophage(+) and prophage(-) backgrounds. Each colored segment indicates the fraction of total observed mutations assigned to a given host function pathway. Pathway distributions differed significantly between ancestral backgrounds (permutation chi-square test, 10,000 label shuffles, $P<0.0001$). As in Figure 3, quinolone resistance pathways (DNA gyrase and efflux pump) were mutated in both backgrounds, but comprised a larger fraction of mutations in prophage(+), but this enrichment was not significant (Monte Carlo permutation test; Fisher's exact test, n.s.).

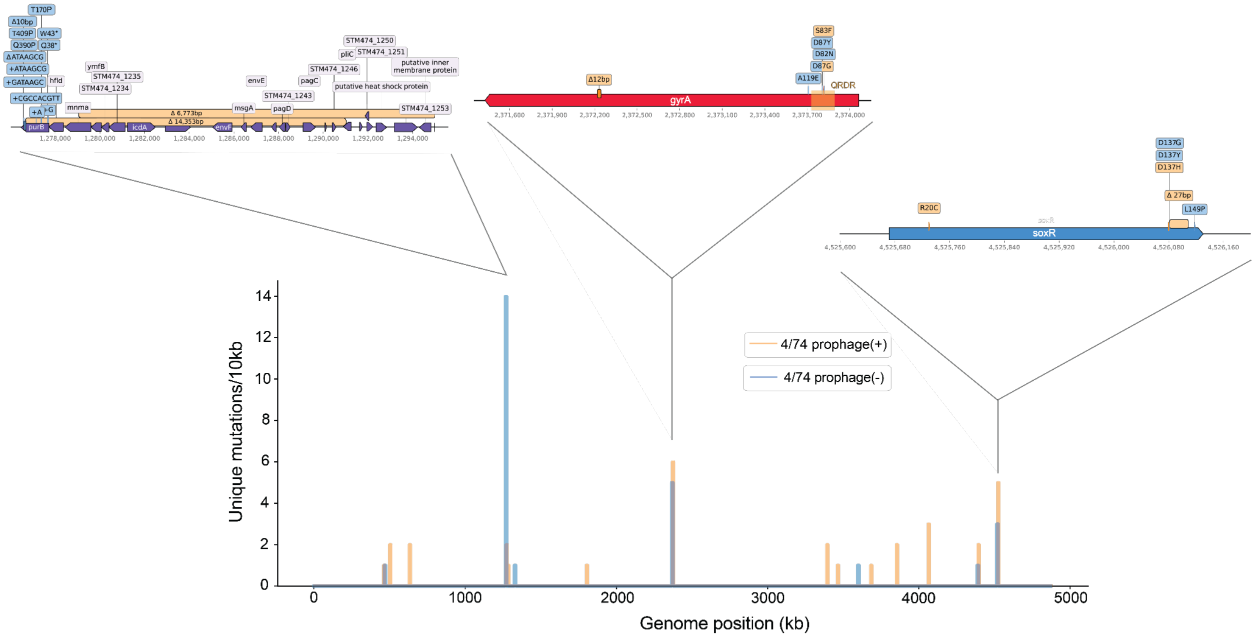


**Supplementary Figure 6 (S6). Distributions of mutated residues differ by prophage carriage in mutation location and type.**

Genomic distribution of unique resistance-associated mutations in prophage(+) and prophage(-) CIP-resistant mutants. Vertical lines within 10 kb genomic bins indicate the presence of one or more mutations. Insets highlight regions of elevated mutational variability, including *purB*, *gyrA*, and *soxR*. Annotated genes are shown as solid arrows with labels indicated on or above (gray box) each feature. Mutations are represented by colored markers corresponding to prophage(+) and prophage(-) mutant backgrounds (orange and blue respectively).

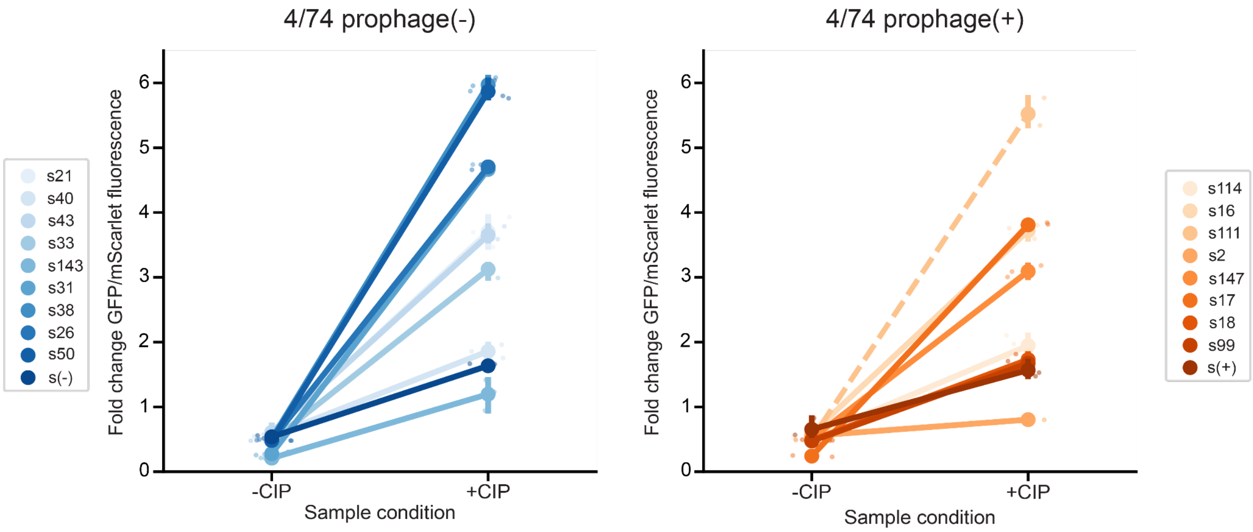


**Supplementary Figure 7 (S7). SOS response induction is dampened in lysogenic mutants, while non-lysogen metabolism mutants maintain high SOS induction.**
Mutant-specific cell-size (mScarlet-i3) normalized precA-GFP expression in the absence (−CIP) and presence (+CIP) of sub-inhibitory ciprofloxacin for prophage(−) (blue) and prophage(+) (orange) resistant mutants. Lines connect measurements from the same mutant across conditions, illustrating changes in SOS induction following ciprofloxacin exposure. Shades represent individual mutants (identities listed in Table S3). Small points indicate biological replicates (n = 3), and large points indicate the mean value for each mutant. Under sub-inhibitory ciprofloxacin treatment, prophage(+) mutants exhibited lower SOS activation and a reduced fold change in GFP expression compared with prophage(−) mutants (Mann–Whitney U test, P = 0.0144). Sample 111 (s111; dashed line) carries a mutation in *clpP* and was considered incompatible with the precA-GFP reporter, as disruption of *clpP* is predicted to impair SOS signaling downstream of recA activation.

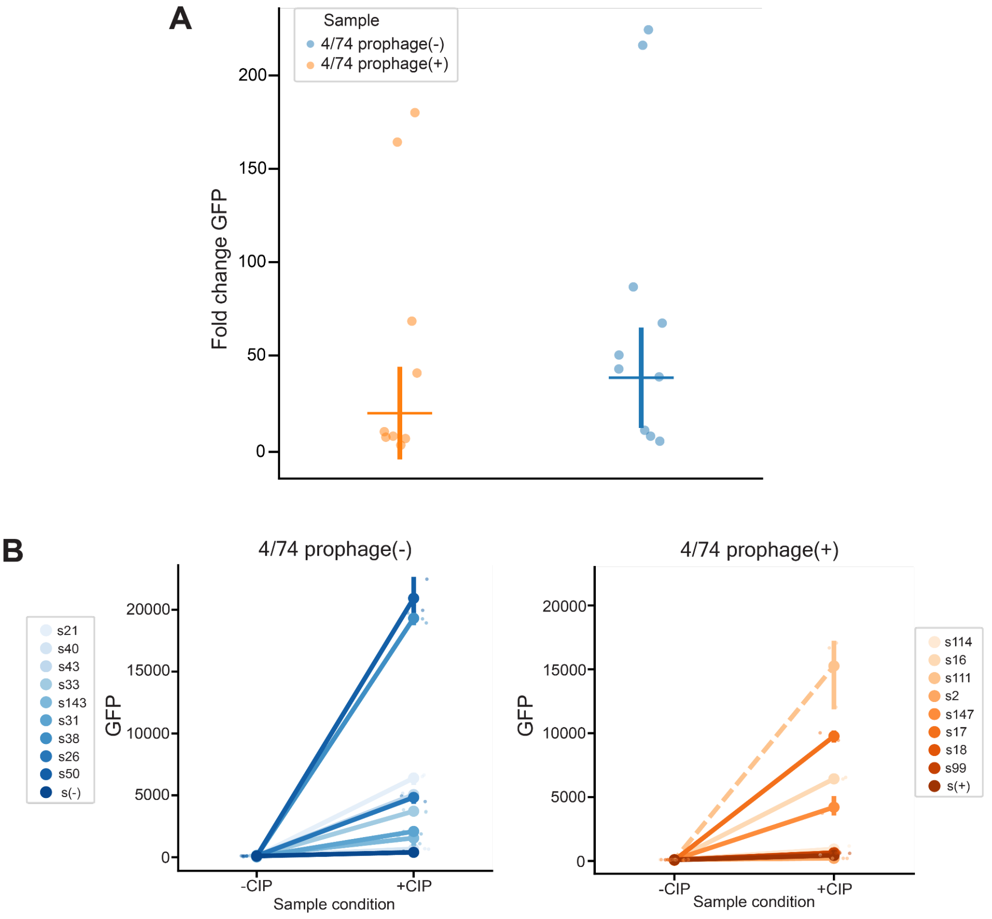


**Supplementary Figure 8 (S8). Dampened SOS response of CIP-resistant lysogens is consistent with or without cell size normalization.**

**A.** Fold change in precA-GFP expression across −CIP and +CIP (sub-inhibitory) conditions for prophage(-) (blue) and prophage(+) (orange) resistant mutants. Each point represents a single mutant's mean GFP expression for 3 biological replicates. Prophage(+) mutants showed significantly lower average induction than prophage(-) mutants, matching the normalized results (Fig. S7, Mann–Whitney U test, $P = 0.03245$). **B.** Mutant-specific precA-GFP expression in the absence (−CIP) and presence (+CIP) of sub-inhibitory ciprofloxacin for prophage(−) (blue) and prophage(+) (orange) resistant mutants. Lines connect measurements from the same mutant across conditions, illustrating changes in SOS induction following ciprofloxacin exposure. Shades represent individual mutants (identities listed in Table S3). Small points indicate biological replicates ($n = 3$), and large points indicate the mean value for each mutant. Under sub-inhibitory ciprofloxacin treatment, prophage(+) mutants exhibited lower SOS activation and a reduced fold change in GFP expression compared with prophage(−) mutants, matching normalized results (Fig. S7). As under normalized conditions, sample 111 (s111; dashed line) carries a mutation in *clpP* and was considered incompatible with the precA-GFP reporter, as disruption of *clpP* is predicted to impair SOS signaling downstream of recA activation.

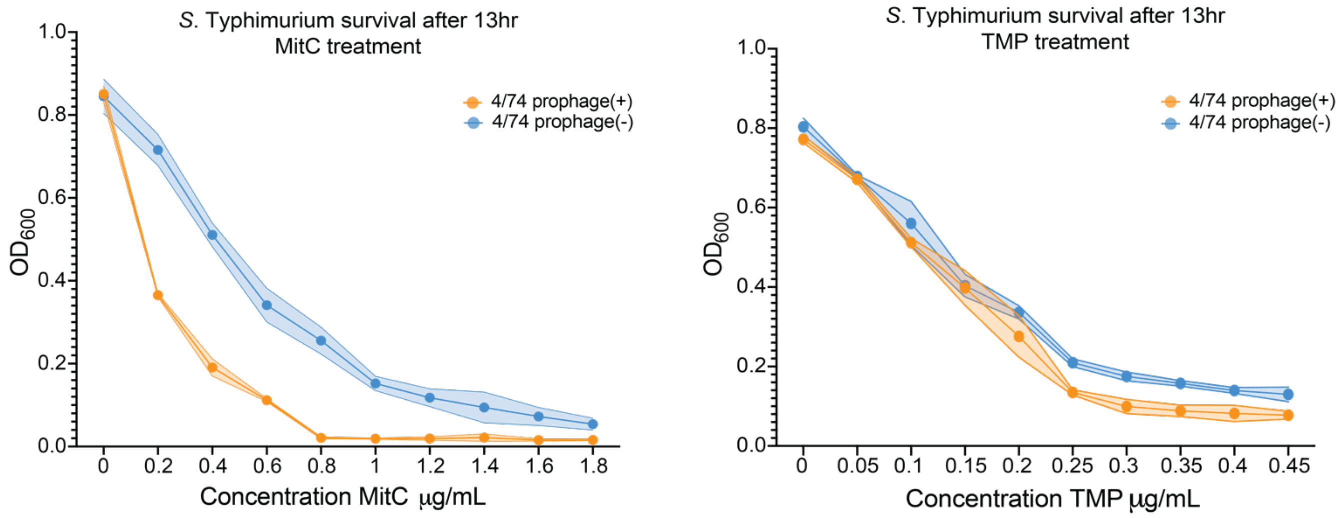


**Supplementary Figure 9 (S9).**

$OD_{600}$ values of prophage(+) and prophage(-) strains showing survival after 13 h growth in increasing concentrations of antibiotic treatment (MitC & TMP, x-axis) as a measure of sensitivity. Points represent mean ± SD of biological replicates. Survival of prophage(+) samples was significantly lower than the non-lysogen at every MitC treatment (multiple *t*-tests with FDR correction, adjusted $P < 0.001$). TMP treatment yielded little to no differences in antibiotic sensitivity across strains. Although significant at high TMP concentrations ($P < 0.01$), these differing values were close to the limit of detection for the plate reader.
